# Red-light is an environmental effector for mutualism between begomovirus and its vector whitefly

**DOI:** 10.1101/2020.07.03.186262

**Authors:** Pingzhi Zhao, Xuan Zhang, Yuqing Gong, Dongqing Xu, Ning Wang, Yanwei Sun, Shu-Sheng Liu, Xing-Wang Deng, Daniel J. Kliebenstein, XuePing Zhou, Rong-Xiang Fang, Jian Ye

## Abstract

Environments such as light condition influence the spread of infectious diseases by affecting insect vector behavior. However, whether and how light affects the host defense which further affect insect preference and performance, remains unclear, nor has been demonstrated how pathogens co-adapt light condition to facilitate vector transmission. We previously showed that begomoviral βC1 inhibits MYC2-mediated jasmonate signaling to establish plant-dependent mutualism with its insect vector. Here we show red-light as an environmental catalyzer to promote mutualism of whitefly-begomovirus by stabilizing βC1, which interacts with PHYTOCHROME-INTERACTING FACTORS (PIFs) transcription factors. PIFs positively control plant defenses against whitefly by directly binding to the promoter of terpene synthase genes and promoting their transcription. Moreover, PIFs integrates light and jasmonate signaling by interaction with MYC2, and co-regulation the transcription of terpene synthase genes. However, begomovirus encoded βC1 inhibit PIFs’ and MYC2’ transcriptional activity via disturbing their dimerization, thereby impairing plant defenses against whitefly-transmitted begomoviruses. Our results thus describe how a viral pathogen hijacks host light signaling to enhance the mutualistic relationship with its insect vector.

**Author summary:** Climate change is driving disease rapidly spread, esp. for global distribution of insect-borne diseases. This paper reports red-light as an environmental factor to promote insect vector olfactory orientation behavior and increase viral disease transmission. Plant virus adapts the supplemental red lighting practice in modern agricultural greenhouse production under protection, therefore enhancing disease spreading globally.

## Introduction

Climate change affect the emergence and spread of vector-borne infectious disease such as malaria, West Nile virus, Zika virus, and viral disease in staple crops via many ways [1, 2]. Rising global temperatures can push disease-carrying insects such as mosquitoes and whiteflies to move into new places that affect the transmission of local viral pathogens [3]. Evidence suggests that crop production is threatened in complex ways by climate changes in the incidence of pests and pathogens [1, 2]. Changed light condition also affects insect vector orientation and therefore feeding behavior. Arthropod-borne viruses (arboviruses) cause diseases in human and crops, and rely on their vectors for transmission and multiplication [4, 5]. The distribution and population size of disease vectors can be heavily affected by local climate and light conditions. Beside of direct effecting fitness of their vectors, plant pathogens confer indirect effects on their vectors often by manipulating the plant defenses against the vector, e.g. volatile chemical components. These volatile substances act as olfactory clues, but also host-finding cues, defensive substances even sex pheromones [6, 7]. Many of insect-borne plant pathogens, e.g. arboviruses of the families *Geminiviridae*, are capable of achieving indirect mutualistic relationships with vectors via their shared host plant [8–10].

To cope with these environmental changes, sessile plants have evolved integrated mechanisms to respond these complex stress conditions, minimizing damages, while conserving valuable resources for growth and reproduction [11–13]. As an energy source and a key environmental factor, light influences plant growth, defense, and even ecological structure [14, 15]. The perception of light signals by phytochrome photoreceptors initiates downstream signaling pathways and regulates numerous plant processes during growth and defense [16]. The *Arabidopsis thaliana* PHYTOCHROME INTERACTING FACTORS (PIFs) are a class of basic helix-loop-helix (bHLH) transcription factors in *Arabidopsis*, which interact with the active photoreceptors to optimize plant growth and development [17–19].

Exogenous light signals integrated with endogenous signals from defense hormones such as jasmonate (JA) and salicylic acid in plant, mediate plant defense responses [14]. These defensive arsenals often produce a blend of ecologically important volatile chemicals such as terpenoids releasing to the environment, and counter the herbivore attack including vectors such as whitefly and aphid [14, 20]. The downstream bHLH transcription factor MYC2 controls the production of some secondary metabolites, which can function as olfactory cues for insects, e.g. terpenoids and glucosinolates [21–24]. Although light is known to regulate plant growth and defense against insects and pathogens [25], how light affects host interaction with herbivore and pathogens has not been described in detail.

Begomovirus, the largest genus of plant viruses and transmitted exclusively by whitefly, have evolved strategies to manipulate JA-regulated plant olfactory cues to promote their mutualism with whitefly vectors [9, 22, 26]. For example, the begomovirus *Tomato yellow leaf curl China virus* (TYLCCNV), which possesses only the DNA-A component with a betasatellite (TYLCCNB), is a whitefly-transmitted begomovirus that results in epidemic diseases in tomato, tobacco and other crops [27–29]. These host plants produce volatile terpenoids as olfactory repellents against whitefly [22, 26, 30]. We have previously shown that the TYLCCNB encoded a βC1 protein suppresses the transcriptional activation-activity of MYC2 by interfering with its dimerization, leading to reduced transcription of *TERPENE SYNTHASE* (*TPS*) genes and terpenoid biosynthesis and anti-herbivory glucosinolates biosynthesis, thereby establishing an indirect mutualistic relationship between the pathogen and the vector [22]. Whether and how climate condition such as light affects this mutualism between begomovirus and whitefly herbivore has not been characterized.

Here, we report red-light as an environmental catalyzer to promote mutualism of whitefly-begomovirus by stabilizing begomovirus-encoded βC1. The family of multiple signaling integrator PIFs is a new key target of the viral βC1 protein. βC1 protein hijacks two kinds of bHLH transcription factors (MYC2 and PIFs) to decrease the transcription of *TPSs* genes that are expected to reduce terpene biosynthesis. Our results show that a begomovirus establishes an indirect mutualistic relationship with whitefly vector by modulating red light and JA signaling-mediated plant defense.

## Results

### Environment red-light is indispensable for betasatellite-encoded βC1 protein to promote host whitefly attraction

To detect whether light affects the natural begomoviral transmission process, we performed whitefly two-choice experiments using *Nicotiana benthamiana* (*Nb*) plants and *Nb* plants infected with TYLCCNV and its associated betasatellite (TYLCCNB) (TA+β) in light and dark conditions (Fig 1A). Consistent with our previous report [22], whiteflies showed a significant preference for TA+β-infected plants to uninfected Nb plants under white light (Fig 1B). Interestingly, the whiteflies did not show preference for TA+β-infected plants under darkness (Fig 1B). We previously demonstrated that βC1 protein encoded by TYLCCNB is involved in host preference of whitefly [22]. More whiteflies were attracted to TA+β-infected plants compared with the βC1 betasatellite mutant virus (TA+mβ)-infected plants under white light, but there were no significant changes of whitefly preference between TA+β-infected plants and TA+mβ-infected plants under darkness (Fig 1B), suggesting that the viral βC1-mediated whitefly preference is light-dependent.

**Fig 1.**
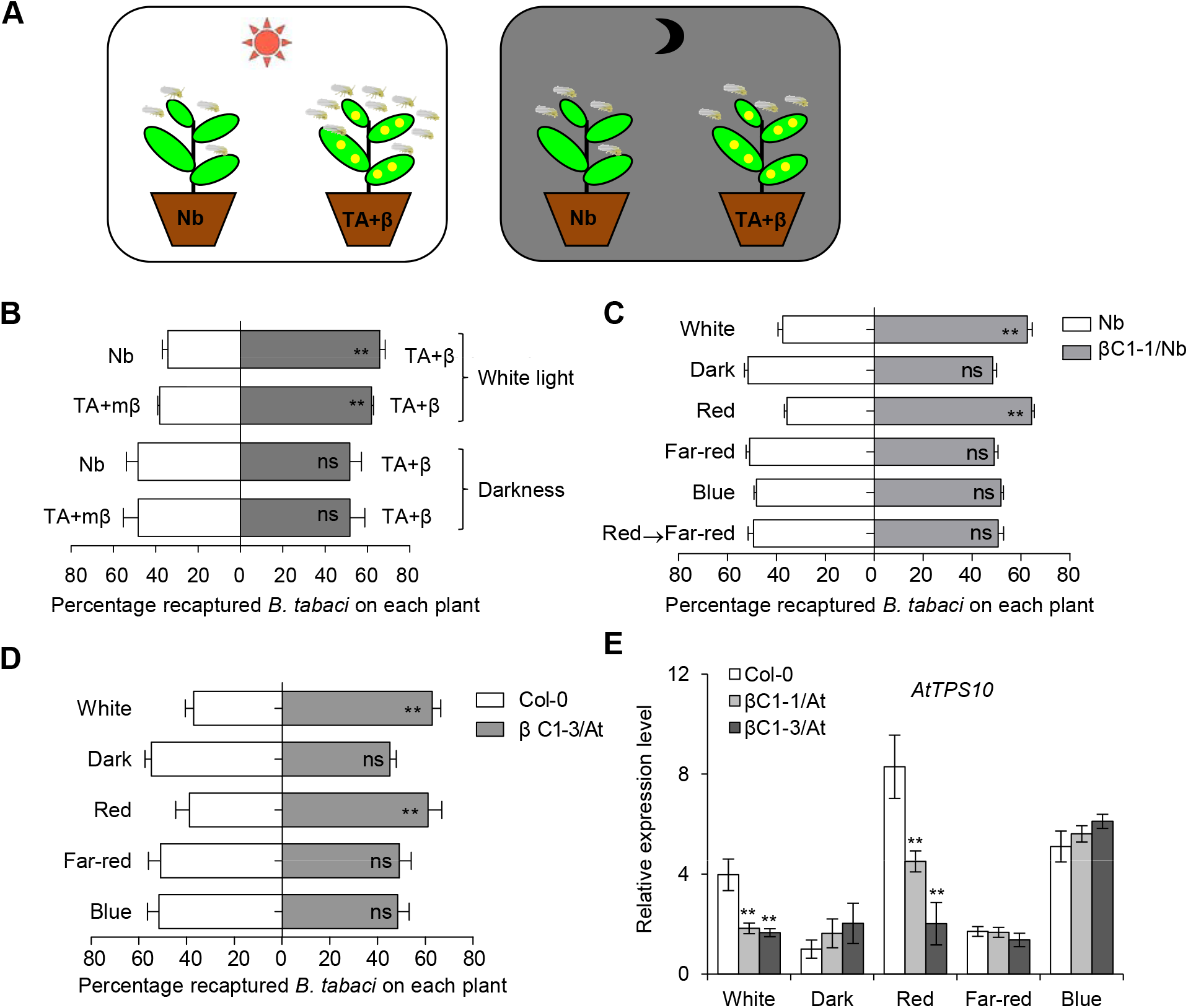
Begomoviral βC1-mediated whitefly preference is light-dependent. **(A)** The schematic diagram of whitefly preference for plant leaf of different ages. Control plants (TA+mβ) or virus-infected plants (TA+β) were exposed to white light (left panel) or dark condition for 12 h at 25℃ (right panel) before whitefly preference experiment. Light intensity was measured using a spectrometer. **(B)** Whitefly preference (as percentage recaptured whiteflies out of 200 released) on uninfected *N. benthamiana* (Nb, mock) and plants infected by TA+β, or on plants infected by TA+β and a mutant βC1 (TA+mβ) in the white light or under darkness. Values are mean + SD (n=6). **(C-D)** Whitefly preference on wild-type Nb and βC1 transgenic Nb plants (βC1-1/Nb) (C) or wild-type Col-0 and βC1 transgenic *Arabidopsis* plants (βC1-3/At) (D) in response to white, dark, red, far-red, and blue light. Plants were placed under darkness for 24 h, followed by a 2 h light exposure and then performed whitefly choice experiments. Red→Far-red indicates that plants were firstly kept in darkness for 24 h, followed by a 2 h red light exposure, and then transferred to far-red light for 2 h. Values are mean + SD (n=6). In B-D, asterisks indicate significant differences between different treatments or lines (**, P< 0.01; ns, no significant differences; the Wilcoxon matched pairs test). **(E)** Relative expression levels of *AtTPS10* in Col-0 and two βC1/At plants (βC1-1/At and βC1-3/At) under different light conditions. Values are mean ± SD (n=3) (*, P< 0.05; **, P< 0.01; Student’s *t*-test). The light was supplied by LED light sources, with irradiance fluency rates of: white (80 μmol m^−2^ sec^−1^), blue (15 μmol m^−2^ sec^−1^), red (20 μmol m^−2^ sec^−1^), and far-red (2 μmol m^−2^ sec^−1^).

We next performed whitefly two-choice assays using *βC1* transgenic Nb plants (βC1/Nb) in various light conditions. Due to the complicity of the tripartite interactions, we applied monochromatic red-light to represent high red: far-red (R:FR) light ratio and also far-red light to represent low R: FR, the latter mimics the poor light condition of plant competition for light. Different monochromatic light sources were used to determine which wavelength of light is essential for whitefly attraction. Whiteflies were more attracted to the βC1/Nb plants compared to wild-type Nb plants only under red light and white light, not in darkness, far-red light and blue light (Fig 1C and S1A Fig). Moreover, red light-induced whitefly attraction from βC1/Nb plants was disrupted by far-red light (Fig 1C and S1A Fig). These whitefly preference results under monochromatic lights agree with the field experiments [31], which encouraged us to design other experiments for explicit mechanisms of light effect on the tripartite interactions. To further confirm these results with another host plant, transgenic *Arabidopsis* plants expressing *βC1* (βC1/At) were used to perform whitefly two-choice assays and the same result as in βC1/Nb plants was observed (Fig 1D). Moreover, wild-type Col-0 plants under red light conferred stronger repellence to whitefly than that under darkness on wild-type Col-0 plants (S2A Fig). These results demonstrate that red light plays a crucial role in whitefly preference for *βC1*-expressing plants.

Hemipterans lack red light photoreceptors, so red light likely cannot directly affect whitefly behaviors [32, 33]. We thus hypothesized that signals from the host plant mediates the red light-induced changes in whitefly preference for plants expressing *βC1*. Our previous work showed that TYLCCNB βC1 contributes to the suppression of JA-regulated terpene biosynthesis and renders virus-infected plant more attractive to its whitefly vector [22]. Therefore, we examined the expression levels of *TPS* genes under various monochromatic light conditions in *Arabidopsis*. Only red light could induce the βC1-mediated suppression of *AtTPS10*, *AtTPS14,* and *AtTPS21* expression (Fig 1E and S1B, S1C Fig). We also found that red light induced higher expression of *AtTPS10* than darkness (S2B Fig). These results revealed that βC1 inhibits the transcription of *TPS* genes in a red light-dependent manner.

### Environment red-light stabilizes βC1 protein in plants

To explore the potential mechanism underlying the interaction between βC1 and plant signaling under various light conditions, we first excluded the possible roles of light on the subcellular localization of βC1 protein or its transcript levels (S3 Fig). We found that the abundance of βC1 was higher under red light than under darkness, far-red light and blue light (Fig 2A). The profile of protein accumulation offered an explanation for the βC1-induced host preference in a red-light dependent manner.

**Fig 2.**
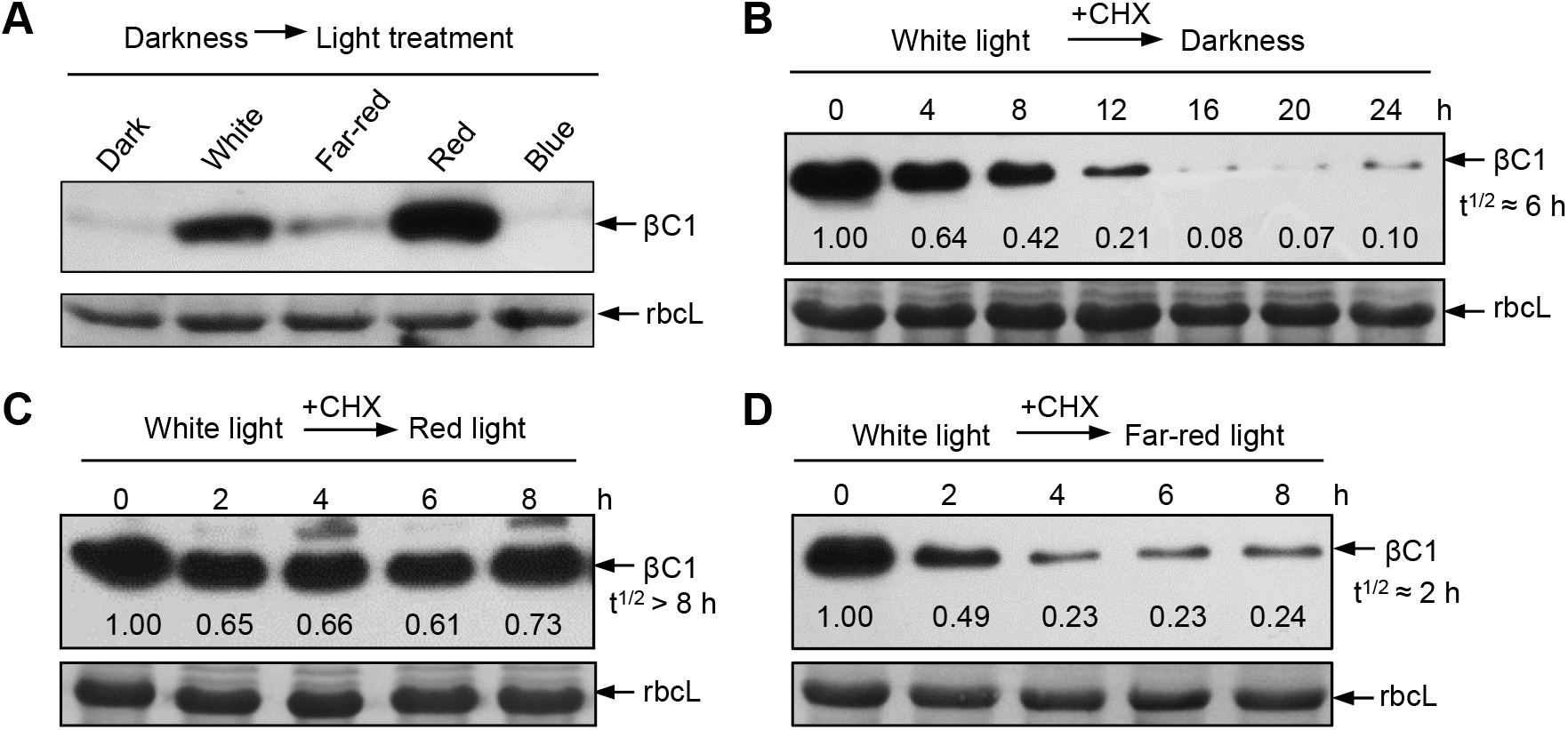
Light-dependent stability of βC1 protein. **(A)** Accumulation of βC1 proteins in Nb plants after different light treatments for 2h. Plants were agroinfiltrated with *35S:myc-βC1*, incubated in the dark for 60 h, and followed by a 2 h light exposure. Samples were detected by immunoblot analysis using anti-myc antibody. Stained membrane bands of the large subunit of Rubisco (rbcL) were used as a loading control. **(B-D)** Degradation of βC1 proteins in response to darkness, red light and far-red light. Nb plants were infiltrated with *A. tumefaciens* cells harboring *35S:myc-βC1*, and incubated in a growth chamber at 25°C with a 12 h light/ 12 h darkness cycle for 60 h. Samples were injected with 100 μM cycloheximide (CHX) and then transferred to darkness (B), exposed to continuous red light (20 μmol m^−2^ sec^−1^) (C), or far-red light (2 μmol m^−2^ sec^−1^) (D), respectively. Samples were collected at the designated times intervals and detected by anti-myc antibody. βC1 protein was quantitated by band intensities in immunoblots using ImageJ software and normalized to individual rbcL level. T^1/2^ indicates half-life of βC1 protein under darkness or various light conditions. Accumulated βC1 protein level at time 0 was set as one.

To further determine the effects of light signals on βC1 stability, Nb plants transiently expressing βC1 were transferred from white light into monochromatic light boxes and sampled at designated time intervals (Fig 2B-D). The accumulation of βC1 sharply decreased under darkness and in far-red light (Fig 2B and 2D). The half-life of βC1 protein was approximately 6 h under darkness and decreased to 2 h under far-red light, suggesting that far-red light signal promotes the degradation of βC1 protein. The protein stability of βC1 protein under red light was much higher than under other monochromatic light and dark condition (Compare Fig 2C with 2B and 2D). Furthermore, we detected the accumulation of βC1 protein in two stable transgenic *Arabidopsis* lines expressing *βC1* (*35S:myc-βC1* #1 and #2). The results show that compared to darkness, white light or red light promotes the stability of βC1 protein in stable transgenic plants (S4 Fig). These results further support a conclusion that red light promotes the stability of βC1.

### βC1 interacts with PIFs

To explore how light signal influences βC1 stability and βC1-induced whitefly attraction, we used a yeast two-hybrid system to screen for βC1 interactors in an *Arabidopsis* cDNA library. This identified AtPIF3, which was first identified to act in the light transduction pathway and later as multiple signaling integrator [34], as a new βC1-targeted host factor. We next confirmed that βC1 interacts with all four of the *Arabidopsis* PIF-quartet proteins (AtPIF1, AtPIF3, AtPIF4, or AtPIF5) by yeast-two hybrid and bimolecular fluorescence complementation (BiFC) assays (Fig 3A and 3B), further co-immunoprecipitation (CoIP) assay confirmed the interaction between βC1 and AtPIF3 *in vivo* (Fig 3C). Taken together, these data suggest that βC1 interacts with PIFs in plants.

**Fig 3.**
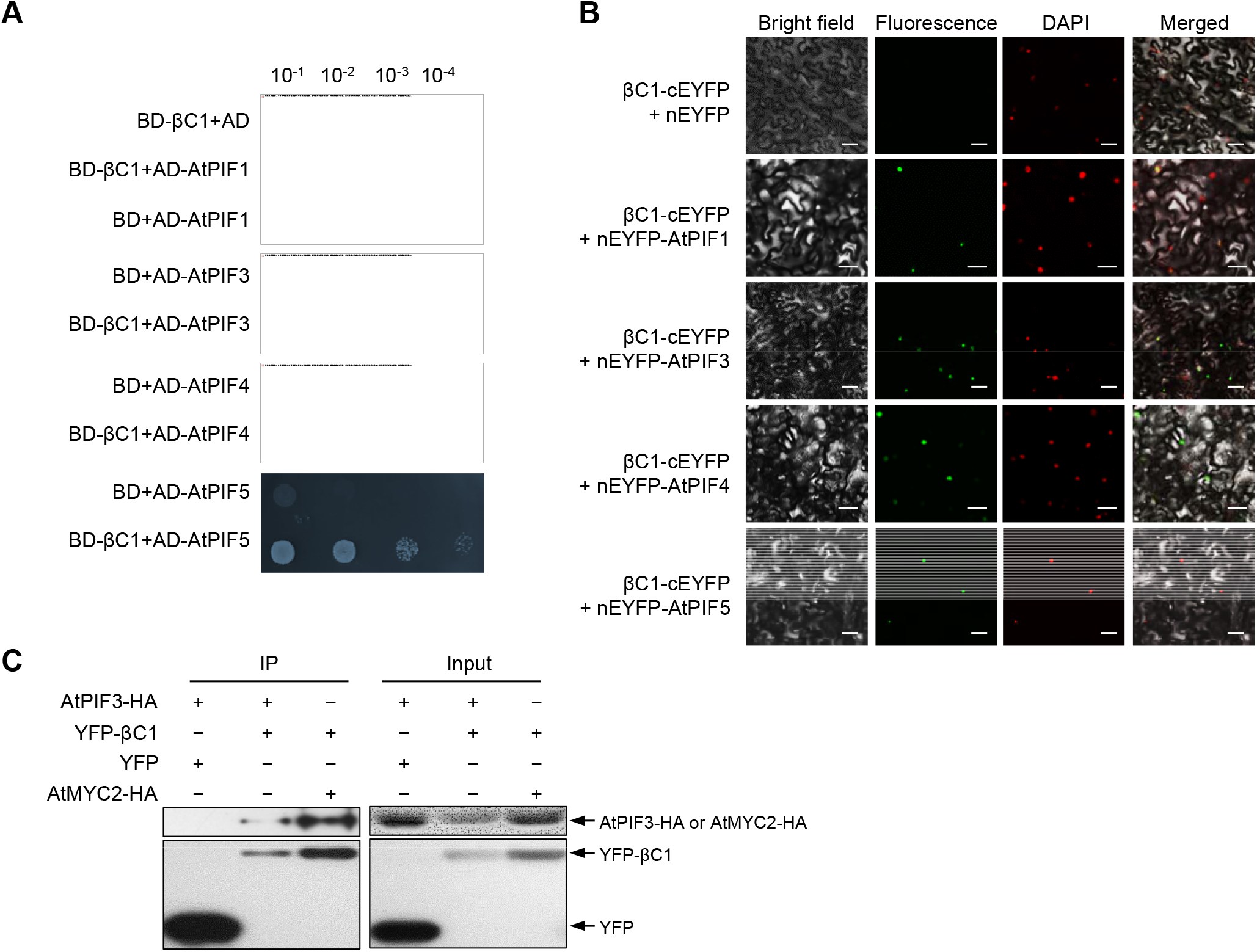
βC1 interacts with phytochrome-interacting factors (PIFs). **(A)** Interaction between βC1 and *Arabidopsis* PIFs (AtPIF1, AtPIF3, AtPIF4 or AtPIF5) in the yeast two-hybrid system. The empty vectors pGAD424 and pGBT9 were used as negative controls. **(B)***In vivo* BiFC analysis of βC1 interaction with *Arabidopsis* PIFs (AtPIF1, AtPIF3, AtPIF4 or AtPIF5). Fluorescence was observed owing to complementation of the βC1-cEYFP fused protein and nEYFP-AtPIFs fused protein. Nuclei of Nb leaf epidermal cells were stained with DAPI. Unfused nEYFP was used as a negative control. Scale bars = 50 μm. **(C)** Co-IP analysis of AtPIF3-HA and YFP-βC1 interaction *in vivo*. YFP was used as a negative control, while AtMYC2-HA was used as a positive control. All of above interaction experiments were performed in normal light condition.

PIFs contain a conserved bHLH domain that binds to DNA and mediates dimerization with other bHLH transcription factors to regulate downstream signaling, and another Active Phytochrome A/B-binding domain that interacts with phyA and phyB to sense upstream signaling [19]. We further showed that βC1 interacts with the bHLH domain of AtPIF3 (S5 Fig), indicating that the interaction between βC1 and PIFs may influence the downstream signaling integrator roles of PIFs in cells.

### The PIFs mediate defense against whitefly in Arabidopsis

To investigate whether PIFs are involved in plant defense against insect vectors, we performed whitefly bioassays using Col-0 and *AtPIF3-*overexpressing (*AtPIF3-OE*) transgenic plants. Whiteflies laid fewer eggs and exhibited slower pupa development on *AtPIF3-OE* transgenic plants than that on Col-0 plants (Fig 4A and 4B). Conversely, they laid more eggs and exhibited faster pupa development on *pifq* (*pif1/3/4/5*) quadruple mutant than that on Col-0 plants (Fig 4C and 4D), an observation is similar to that with *βC1*-expressing *Arabidopsis* plants [22]. These data suggest that PIFs are involved in plant defense against whitefly vector.

**Fig 4.**
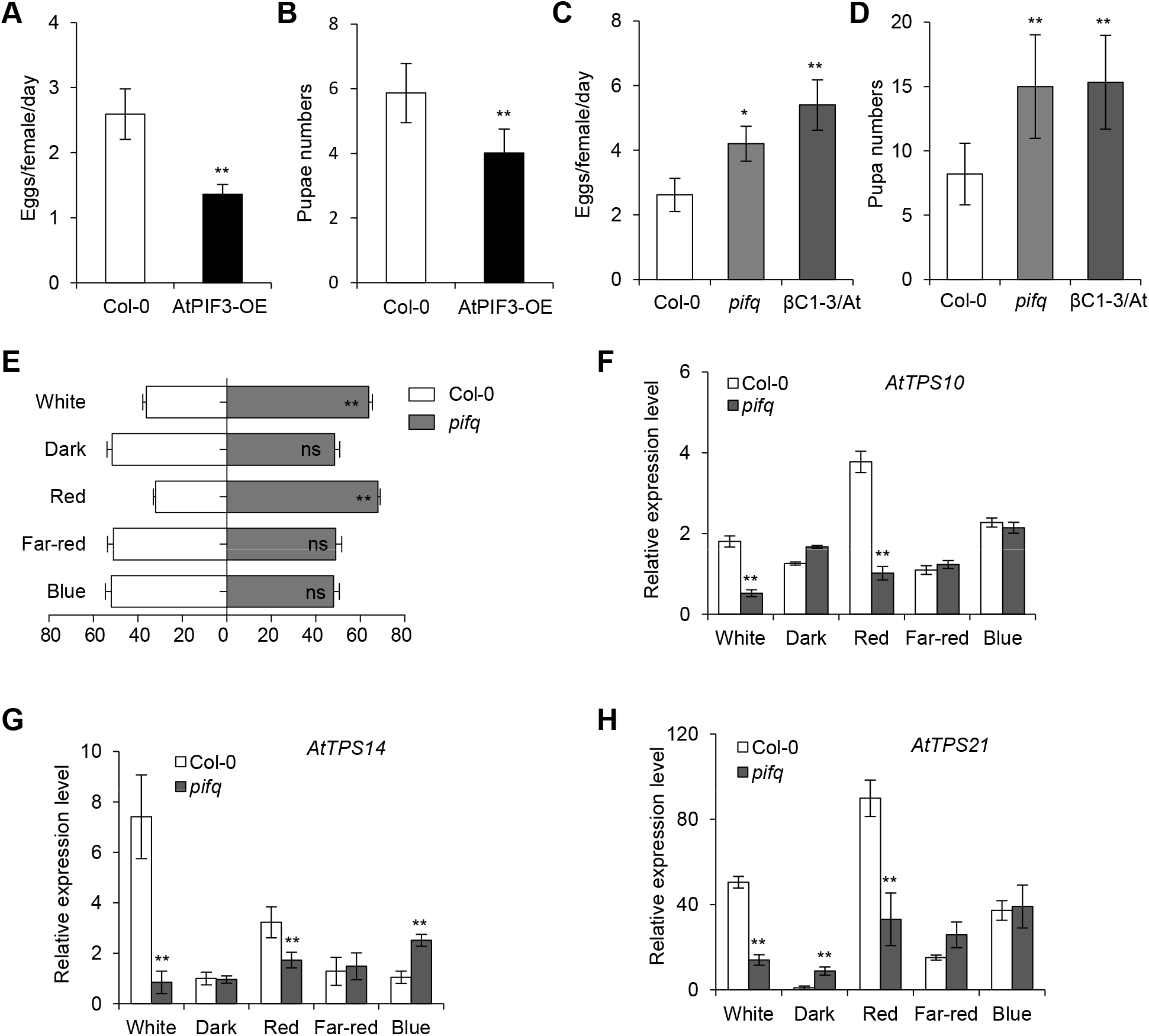
*Arabidopsis* PIFs confer tolerance to whitefly vector. **(A)** Number of eggs laid per female whitefly per day on Col-0 and *AtPIF3*-overexpressing (AtPIF3-OE) transgenic plants. **(B)** Pupa numbers of whiteflies on Col-0 and AtPIF3-OE transgenic plants. **(C)** Number of eggs laid per female whitefly per day on Col-0, *pifq* or βC1-3/At plants. **(D)** Pupa numbers of whiteflies on Col-0, *pifq* or βC1-3/At plants. In figure A-D, values are mean ± SD (n=8). Asterisks indicate significant differences of whitefly performance between Col-0 and mutant plants (*, P< 0.05; **, P< 0.01; Student’s *t*-test). **(E)** Whitefly preference on Col-0 and *pifq* mutant plants in response to white, dark, red, far-red, and blue light. The plants were placed in darkness for 24 h prior to the 2 h different light treatments. Values are mean + SD (n=6) (**, P< 0.01; ns, no significant differences; the Wilcoxon matched pairs test). **(F-H)** Relative expression levels of *AtTPS10* (F)*, AtTPS14* (G), and *AtTPS21* (H) in Col-0 and *pifq* mutant plants after a 2 h treatment of different lights. Values are mean ± SD (n=3) (**, P< 0.01; Student’s *t*-test).

Since TYLCCNB βC1 involved in whitefly preference under white light (Fig 1) and interacts with plant PIFs (Fig 3), we performed whitefly two-choice assays to examine whether the PIFs have the same effects on whitefly preference as βC1. Consistent with the results with *βC1*-expressing *Arabidopsis* plants, the *pifq* quadruple mutants were higher attractive to whiteflies than Col-0 plants under white or red light (Fig 4E). The transcriptional levels of *TPS* genes (*AtTPS10*, *AtTPS14* and *AtTPS21*) were significantly repressed in the *pifq* mutant compared to those in Col-0 plants under white or red light (Fig 4F-H). Taken together, these results imply that transgenic expression of *βC1* partially mimics the *pifq* mutant, which hinders the plant terpene-based resistance to whitefly.

### βC1 suppresses PIFs activity by interfering with its dimerization

PIFs are bHLH transcription factors that directly regulate gene expression by binding to a core G-box motif (CACGTG) and G-box-like motif (CANNTG) [34, 35]. We wonder whether PIFs directly regulate the expression of *TPS* genes and involve in the terpene-mediated whitefly defense response. There are five G-box-like elements (CANNTG) in the promoter of *AtTPS10*, distributed in three regions (Fig 5A). We performed a chromatin immunoprecipitation (ChIP) assay using *AtPIF3-OE* plants. Quantitative PCR analysis showed that region II (one G-box-like motif 0.7 kb upstream of the transcription start site) of *AtTPS10* was significantly enriched in *AtPIF3-OE* lines relative to Col-0 plants (Fig 5B). These data indicate that AtPIF3 directly binds to the promoter of *AtTPS10* and regulates its expression in *Arabidopsis*.

PIF3 activates downstream gene expression by forming homodimers and heterodimers with other PIF-related bHLH transcription factors [19]. The interaction between βC1 and the bHLH domain of AtPIF3 (S5 Fig) raised the possibility that βC1 competes with the bHLH domain to interfere with AtPIFs dimerization. A modified BiFC assay was used to test this hypothesis. In cells co-expressing βC1, the interaction signal strength of AtPIF3-AtPIF3 or AtPIF3-AtPIF4 decreased to approximately half of its original intensity (Fig 5C-E), suggesting that βC1 may interfere with PIF dimerization. Moreover, *in vitro* competitive pull-down assays showed that βC1 interferes with homodimerization of AtPIF4-AtPIF4 and heterodimerization of AtPIF3-AtPIF4 (Fig 5F and 5G).

**Fig 5.**
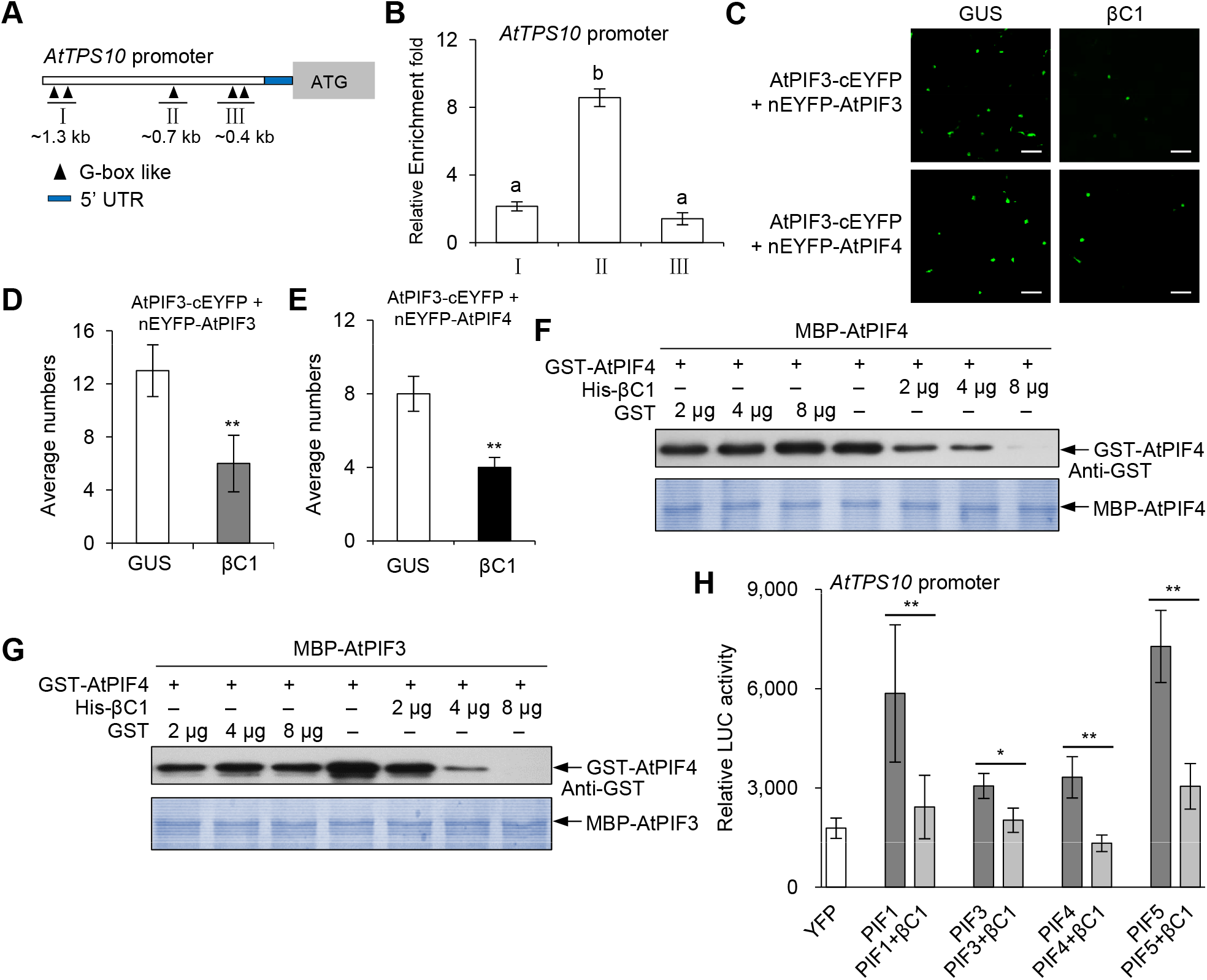
βC1 suppresses transcriptional activity of PIFs by inhibiting its dimerization. **(A)** Schematic diagram of *AtTPS10* promoter. The black triangles represent G-box like motifs. A fragment of the three lines (I, II and III), as indicated by the triangles was amplified in ChIP assay. The end positions of each fragment (kb) relative to the transcription start site are indicated below. UTR, untranslated region. **(B)** Fold enrichment of YFP-AtPIF3 associated with each of the three DNA fragments (I, II and III) of *AtTPS10* promoter in ChIP assay. Values are mean ± SD (n=4). The same letters above the bars indicate lack of significant difference at the 0.05 level by Duncan’s multiple range test. **(C)** Modified BiFC competition assays. The EYFP fluorescence was detected after co-expression of GUS + AtPIF3-cEYFP + nEYFP-AtPIF3 (GUS), βC1 + AtPIF3-cEYFP + nEYFP-AtPIF3 (βC1), or GUS + AtPIF3-cEYFP + nEYFP-AtPIF4, βC1 + AtPIF3-cEYFP + nEYFP-AtPIF4. Scale bars = 50 μm. **(D-E)** Average numbers of EYFP fluorescence show effects of βC1 on the formation of AtPIF3-AtPIF3 homodimers (D) and AtPIF3-AtPIF4 heterodimers (E). Values are mean ± SD (n=8) (**, P< 0.01; Student’s *t*-test). **(F-G)** GST pull-down protein competition assays. The indicated protein amount of His-βC1 or GST was mixed with 2 μg of GST-AtPIF4 and pulled down by 2 μg of MBP-AtPIF4 ( F) or 2 μg of MBP-AtPIF3 ( G). Immunoblots were performed using anti-GST antibody to detect the associated proteins. Membranes were stained with Coomassie brilliant blue to monitor input protein amount. **(H)** Effects of βC1 on transcriptional activity of each AtPIFs (AtPIF1, AtPIF3, AtPIF4, or AtPIF5) on *AtTPS10* promoter under white light. *AtTPS10* promoter: *luciferase* (*LUC*) was used as a reporter construct. YFP, YFP-AtPIFs, and YFP-βC1 were used as effector constructs. Values are mean ± SD (n=8) (*, P< 0.05; **, P< 0.01; Student’s *t*-test).

Next, we examined whether βC1 affects the trans-activity of PIFs via a construct containing the *AtTPS10* promoter with *luciferase* (*LUC*) as a reporter, and YFP-AtPIFs (AtPIF1, AtPIF3, AtPIF4, or AtPIF5) as effectors. *AtTPS10* promoter: *LUC* was transiently expressed with the indicated effector plus βC1 in Nb leaf cells. Fig 5H shows that each of AtPIFs (AtPIF1, AtPIF3, AtPIF4, and AtPIF5) significantly increased the LUC activity, whereas βC1 decreased AtPIFs-induced LUC activity at different degrees (Fig 5H). Taken together, these results indicate that βC1 attenuates the trans-activity of AtPIFs in promoting *AtTPS10* transcription by inhibiting PIF dimerization.

### Light and JA signals coordinately regulate host preference of whitefly

PIF-quartet integrates signals from multiple signaling pathways, including light and JA signals, to respond to the diverse stresses and developmental processes [19, 34, 36]. Previous study has reported that AtPIF4 interacts with AtMYC2, and JA inhibits the function of PIF4 partially through MYC2 in *Arabidopsis* [37]. To confirm that JA and light signaling work cooperatively to regulate plant defense against whitefly, we firstly investigated whether MYC2 associates with PIFs in plants. BiFC assays showed that AtMYC2 interacts with AtPIF3 and AtPIF4 (S6 Fig). Additionally, we generated a *pifq/myc2-1* mutant by crossing the *pifq* mutant with the *myc2-1* mutant. The transcriptional levels of *AtTPS10* in the *pifq/myc2-1* mutant were additively reduced compared to the parental lines under red light conditions (Fig 6A). The results suggest that AtPIFs and AtMYC2 coordinately regulate the expression of *AtTPS10*. Since the individual AtPIF4 or AtMYC2 could directly bind and promote *AtTPS10* expression, we next tested whether the AtPIF4-AtMYC2 interaction has synergetic effect on downstream genes expression regulation. Unexpected, we found that the heterdimerazation of AtPIF4-AtMYC2 in fact even reduces the transactivation activity when co-expressed with AtPIF4 and AtMYC2 compared to AtMYC2 alone under white light (Fig 6B), indicating an antagonistic effect of heterodimer formation of AtPIF4-AtMYC2 on expression of *AtTPS10*.

**Fig 6.**
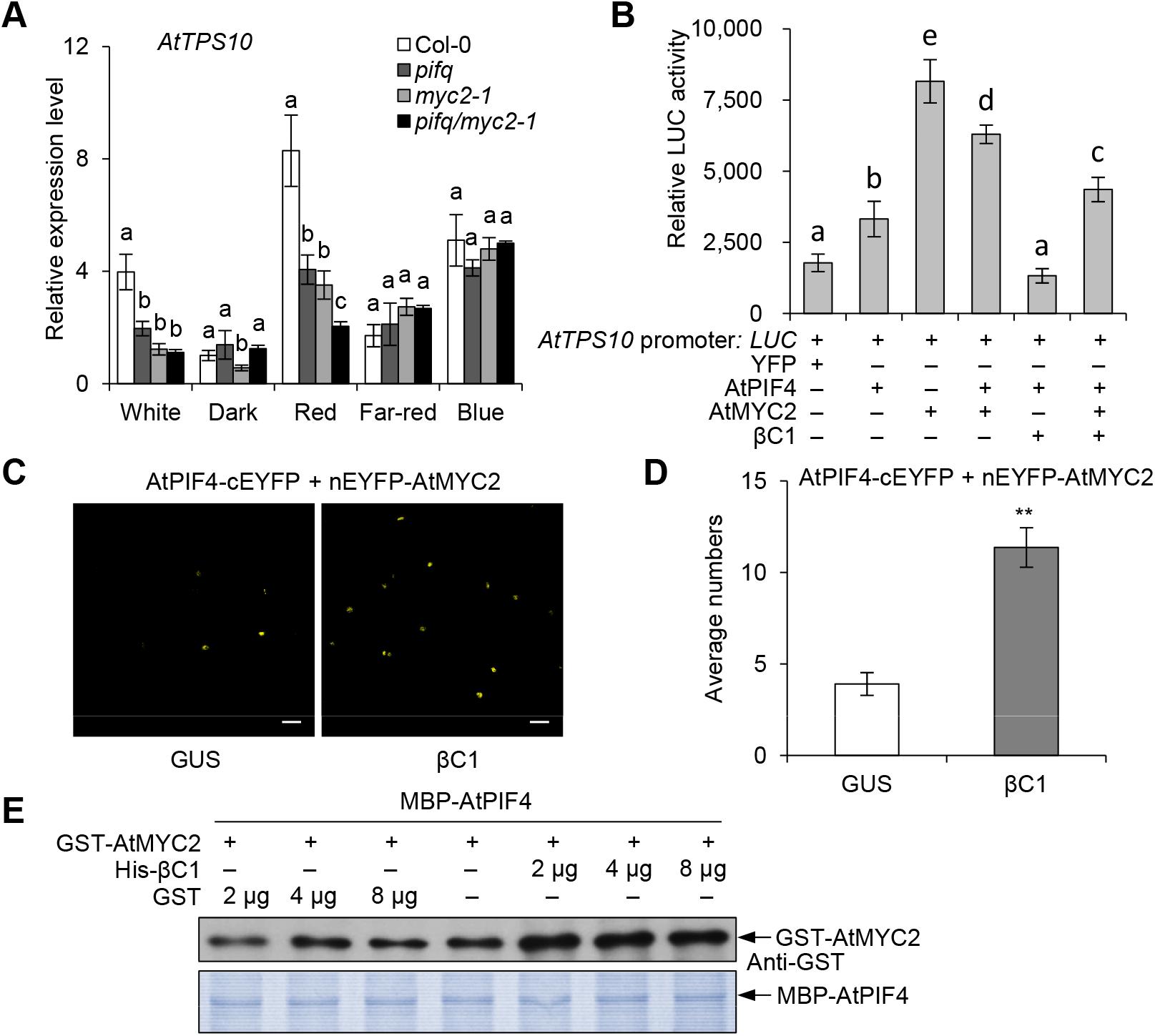
Arabidopsis PIFs and MYC2 transcription factors synergistically regulate *AtTPS10* transcription. **(A)** Relative expression levels of *AtTPS10* in Col-0, *pifq*, *myc2-1* and *pifq/myc2-1* mutant plants after a 2 h treatment with different light. Values are mean ± SD (n=3). **(B)** Effects of βC1 on trans-activation activity of AtPIF4 or AtMYC2 on *AtTPS10* promoter under white light. *AtTPS10* Promoter: *LUC* was used as a reporter construct. YFP, AtPIF4, AtMYC2 and βC1 were used as effector constructs. Values are mean ± SD (n=8). In A and B, the same letters above the bar indicate lack of significant differences at the 0.05 level in Duncan’s multiple range test. **(C)** Modified BiFC competition assays. The EYFP fluorescences were detected using co-expression of AtPIF4-cEYFP + nEYFP-AtMYC2 with or without βC1 under normal light. Scale bars = 50 μm. **(D)** Effects of βC1 on the interaction between AtPIF4 and AtMYC2. Values are mean ± SD (n=8) (**, P< 0.01; Student’s *t*-test). **(E)** Protein competition pull-down assay. The indicated protein amount of His-βC1 or GST was mixed with 2 μg of GST-AtMYC2 and pulled down by 2 μg of MBP-AtPIF4. The associated proteins were detected by immunoblots using anti-GST antibody.

Next we tested the effect of viral βC1 on the AtPIF4-AtMYC2 interaction and found that the interaction signal of AtPIF4-AtMYC2 was increased by two-fold when co-expressed with βC1, but not with β-glucuronidase (GUS) (Fig 6C and 6D), suggesting that βC1 function as a linker between two bHLH transcription factors AtPIF4 and AtMYC2. Competitive pull-down assay also supported the idea that βC1 indeed bridges the interaction of AtPIF4-AtMYC2 (Fig 6E). One hypothesis was then raised that the self-interaction of AtPIFs or AtMYCs promotes their transcriptional activity, but the formation of heterodimer of AtPIF4-AtMYC2 inhibits the MYC2 transcriptional activity. Once plants are infected by begomovirus, the linker-βC1 even exacerbates their activities. For that end, we coexpressed βC1 and found that βC1 could dampen the activator activities either by single AtPIF4 or AtMYC2 or coexpression of these two bHLH transcription factors (Fig 6B).

To further explore the function of JA signals in βC1- or PIFs-mediated whitefly host preference, we performed whitefly two-choice assays using *βC1*-expressing and *pifq* mutant plants with MeJA treatment in darkness. The loss of whitefly preference for βC1/At and *pifq* mutant plants under darkness was rescued by MeJA application (Fig 7A and 7B). Accordingly, the expression levels of *AtTPS10*, *AtTPS14* and *AtTPS21* in two βC1/At lines and *pifq* plants were also dramatically decreased by MeJA under darkness (Fig 7C-H). These results demonstrate that JA and light signals integrally modify begomovirus-whitefly mutualism.

**Fig 7.**
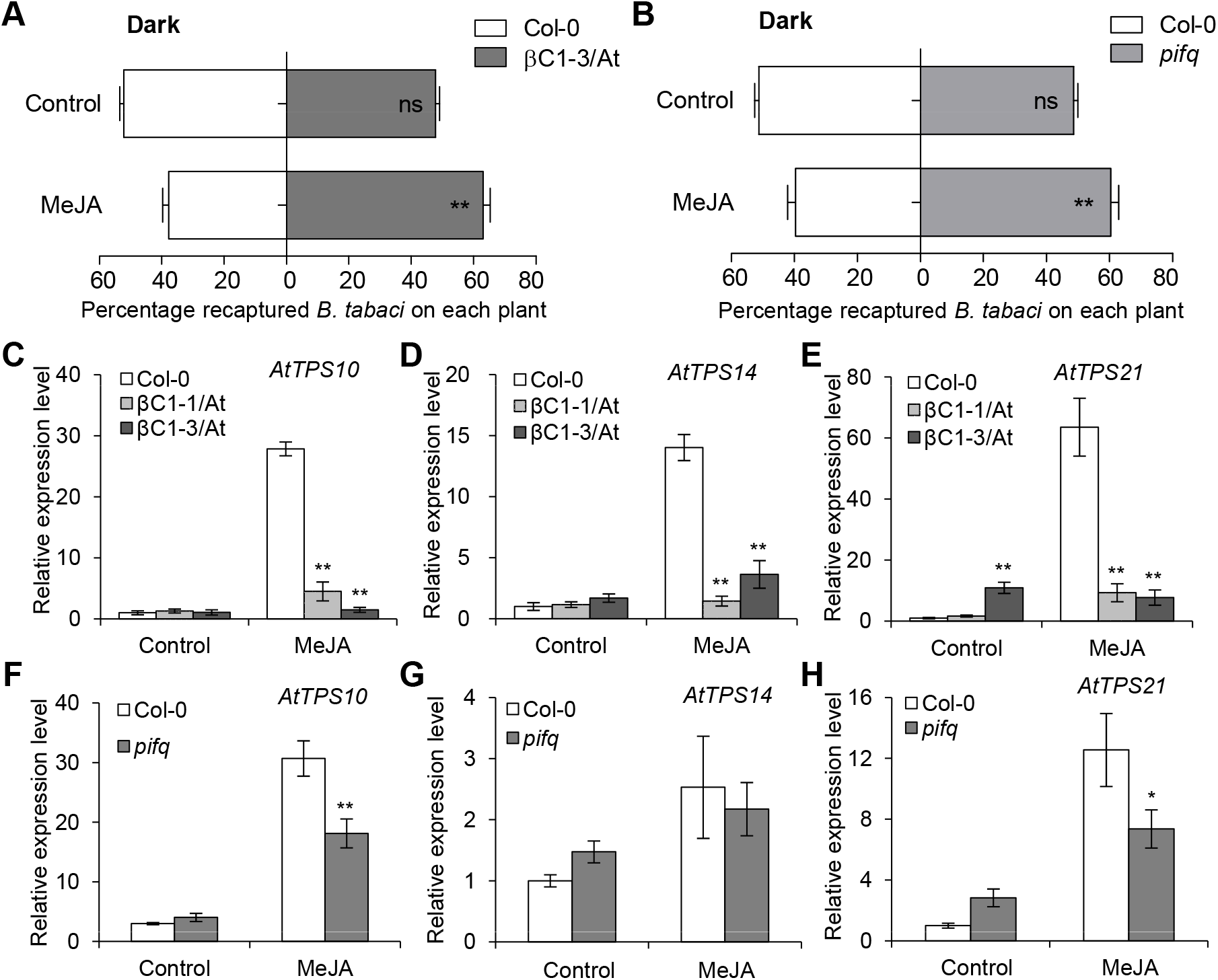
Light and JA signals synergistically regulate whitefly host preference. (A-B) Whitefly preference on Col-0 and βC1-3/At plants (A) or Col-0 and *pifq* mutant plants (B) with or without MeJA treatment under darkness. Values are mean ± SD (n=6) (**, P< 0.01; ns, no significant differences; the Wilcoxon matched pairs test). (**C-E)** Relative expression levels of *AtTPS10* (C), *AtTPS14* (D), and *AtTPS21* (E) in Col-0 and two βC1 transgenic *Arabidopsis* lines with or without MeJA treatment under darkness. Values are mean ± SD (n=3) (**, P< 0.01; Student’s *t*-test). (**F-H)** Relative expression levels of *AtTPS10* (F), *AtTPS14* (G), and *AtTPS21* (H) in Col-0 and *pifq* mutant with or without MeJA treatment under darkness. Values are mean ± SD (n=3) (*, P< 0.05; **, P< 0.01; Student’s *t*-test).

## Discussion

As the Earth warms, we need to be able to predict what conditions will be at risk for infectious diseases because prevention is always superior to reaction. The disease triangle, pathogen-host-environment, is used to understand how disease epidemics can be predicted, restricted or controlled [38]. Evidence is increasing suggesting that environmental factors including light are important mediators of plant defenses during plant-pathogen interactions [14, 34]. However, the ability of plant pathogens in using effectors to disturb or co-opt host light signaling to promote infection has not been well explored. Plant defense signals function as players or pawns in plant-virus-vector interactions [39], PIFs are key signal integrators in regulating plant growth and development [16, 34]. Here, we provide evidence showing that PIFs act as direct positive regulators in plant defense against whitefly vector. First, the *AtPIF3*-overexpression confers enhanced *Arabidopsis* resistance to whitefly, and PIFs deficiency in *pifq* mutant promotes whitefly performance in *Arabidopsis* (Fig 4A-4D). Second, the *pifq* quadruple mutants attract more whiteflies than Col-0 plants under white or red light (Fig 4E). Therefore, PIF is not only directly involved in plant development, but also involved in resistance against vector insects. However, begomoviral βC1 protein performs a successful counter-defense by hijacking PIFs proteins. On the one hand, βC1 interacts with PIFs and suppresses trans-activity of PIFs by interfering with its dimerization (Fig 3 and Fig 5); on the other hand, βC1 utilizes whitefly vector to decrease the *PIFs* transcription induced by begomovirus in host plants (S7 Fig). Consequently, begomovirus suppresses PIFs-mediated plant defense to enhance vector transmission.

Most of plant arboviruses attract their insect vectors by modulating plant host-insect vector specific recognition. Light modulates communications of plant-insect through a combination of olfactory and visual cues comprehensively [14, 40]. Similarly as our current results, red light seems essential for a terpenoid volatile based-attraction to Huanglongbing host plant for the vector insect Asian Citrus Psyllid, which transmits the casual bacterial pathogen *Candidatus Liberibacter* [41]. MYC2 and its homologs have been characterized as a few known regulators in terpene biosynthesis mainly during day time [21–23], since it stabilizes by light but destabilizes in darkness [42]. A recent study demonstrated that the MYC2 protein in JA signaling pathways interacts with PIF4 [37]. Here, we likewise show that AtPIF3 and AtPIF4 all interact with AtMYC2, and *AtTPS10* expression was significantly reduced in *pifq/myc2-1* quintuple mutant compared with parental single *pifq* or *myc2-1* mutant (Fig 6A). Meanwhile transcript accumulation of *TPS* genes (*AtTPS10*, *AtTPS14* and *AtTPS21*) does not show a circadian rhythm in *Arabidopsis* (S8A Fig). The expression of PIFs and MYCs was complemented and balanced regulation (S8B Fig and S8C Fig). These results suggest that PIFs and MYC2 synergistically regulate terpene biosynthesis in two paralleled pathways. Furthermore, the mechanism of PIFs-regulated *TPS* genes expression in dark is complementary to the MYC2-regulated counterpart in light. Since PIFs are much stable in the night, PIFs may control the ecological interactions of plant-insects in night by regulating the chemical communication, esp. for these night blooming plants and behaviors of nighttime feeding insects [43]. In addition, PIF-like genes are highly conserved and they have been existed before the water-to-land transition of plants [16, 44]. It will be of interest to examine possible defensive roles in PIF homologs in other plants. Our findings indicate prospects for biotechnological improvement of crops to improve yield and immunity simultaneously through editing and regulation of *PIFs* genes.

Under red light, PIFs levels/activities are expected to be low in wild-type plants, because PIFs are inactivated by phyB. phyB is the predominant photoreceptor regulating photomorphogenic responses to red light, while phyA is the primary photoreceptor responsible for perceiving far-red light [45, 46]. Our results show that the accumulation of βC1 protein was higher in red light than that in far-red light (Fig 2A). Interestingly, when we treated the Nb plants transiently expressed myc-βC1 protein with continuous red light and far-red light, the βC1 protein was accumulated in red light, but decreased in far-red light (S9 Fig). When the plants in red light again, βC1 protein accumulation has no obvious changes, but reduced again when treated with far-red light (S9 Fig). These results further proved that red light could maintain the stability of βC1 protein, but far-red light promote the degradation of βC1 protein, which imply that the photoreceptors phyB or phyA might involve in regulation of βC1 stability. This hypothesis needs further research.

Modern anti-arbovirus strategy includes anti-insect netting to disrupt disease transmission. Also more and more countries have adapted greenhouse crop production under protected condition in the past decades. In northern countries this practice often relies heavily on supplemental lighting for year-round yield and product quality. Among the different spectra used in supplemental lighting, red light is often considered the most efficient [2]. It seems like that begomovirus could adapt these serial artificial environmental changes by evolving new role of a known virulence factor to hijcak host internal light signaling. Plant viruses have a small genome in which the encoding proteins especially the virulence factors are frequently multifunctional. βC1 proteins are multifunctional and has many host targets for its pathogenesis [29, 47], many of which may impact plant-virus and plant-whitefly interactions. It is necessary to further dissect whether and how other targets of βC1 are also involved in this light-dependent virus pathogenicity in the future. Meanwhile, the data collected here and conclusion we made is based on well-controlled monochromatic light conditions. When extrapolating to natural and agricultural field conditions, it should seriously take into account the real light quality within dense stands in the begomovirus-whitefly-plant tripartite interactions. Nevertheless, our data here is significant for understanding of the tripartite interactions and also for arbovirus disease controlling, esp. begomoviral βC1 is adapted to red-light condition, which represents a good light quality, to suppress phytohormone-regulated terpene biosynthesis to attract whitefly insect.

The results in this study can be best summarized by the working model presented in S10 Fig. In this model, homodimerized PIFs or MYC2 binds to the promoter regions of *TPS* genes, resulting in increased *TPSs* transcript levels and terpene biosynthesis. Thus red-light signal and JA signal fine-tune transcription of *TPS* genes to contribute to resistance to whiteflies in uninfected plants (S10A Fig). In begomovirus-infected plants, βC1 inhibits transcriptional activity of PIFs and MYC2 by interfering with their homodimerization and promoting AtPIFs-AtMYC2 heterodimerization. Finally, the decreased terpene synthesis and in turn enhanced whitefly performance increase the probability of pathogen transmission (S10B Fig).

## Materials and Methods

### Plant materials and growth conditions

Wild-type or transgenic *Nicotiana benthamiana* plants carrying *35S:βC1* have been reported previously [22, 48]. *N. benthamiana* plants grew in an insect-free growth chamber at 25°C with 12 h light/12 h darkness cycle. *Arabidopsis thaliana* wild-type Col-0, *pifq* (*pif1/3/4/5*) [49], *myc2-1* mutant [50], and βC1/At [22] were used in the study. Quintuple *pifq/myc2-1* mutant was generated by crossing the corresponding parental single *myc2-1* and quadruple *pifq* homozygous lines. The construct expressing *35S:YFP-AtPIF3* was transformed into Col-0 plants, and generated *AtPIF3*-overexpressing lines (*AtPIF3-OE*). Sterilized seeds were incubated on Murashige and Skoog medium at 4°C for 3 d before being transferred to a growth chamber (22°C with 10 h of light/14 h of darkness cycle).

### Plant treatments

For whitefly two-choice assays and *Arabidopsis TPSs* expression analysis, plants were placed in darkness for 24 h, followed by a 2-h light exposure for two-choice assays. White light, blue light, red light, and far-red light were supplied by LED light sources, the irradiance fluency rates was, white light (80 μmol m^−2^ sec^−1^), blue light (15 μmol m^−2^ sec^−1^), red light (20 μmol m^−2^ sec^−1^), and far-red light (2 μmol m^−2^ sec^−1^). Light intensity was measured with an OHSP-350C illumination spectrum analyzer.

For phytohormones treatments, methyl jasmonate (MeJA) was used to mimic whitefly infestation in *N. bentheamina* and *Arabidopsis* [22]. Three week-old *Arabidopsis* were sprayed with 100 μM MeJA containing 0.01% (v/v) Tween 20. Plant samples were collected at 6 h following treatment. Control plants were treated with 0.01% (v/v) Tween 20 in parallel for the same time period.

### Virus inoculation

*N. benthamiana* plants with four to six true leaves were infiltrated with *Agrobacterium tumefaciens* carrying TYLCCNV and betasatellite DNAβ (isolate Y10) as described previously [22, 51]. Infiltration with buffer or TYLCCNV plus a mutant betasatellite DNA with a βC1 mutation (TA+mβ) was used as a control [51].

### Whitefly bioassays

Whiteflies were collected in the field in Chaoyang District, Beijing, China and were identified as *Bemisia tabaci* MEAM1, B biotype (mtCOI, GenBank accession number MF579701). The whitefly population was maintained in a growth chamber (25°C, 65% RH) on cotton with a 12 h-light/12 h-dark light cycle.

The whitefly two-choice experiments were performed as described previously [22]. Two plants of selected genotypes with similar size and leaf numbers were firstly kept in darkness for 24 h, and then exposed to specific light for 2 h, and finally placed in an insect cage (30*30*30 cm) with the same light condition. Two hundred adult whiteflies were captured, and then released from the middle of the two plants. After 20 min, the whiteflies settled on each plant were recaptured and the number on each plant was recorded. Six biological replicates were conducted in this experiment.

For whitefly oviposition experiment, three female and three male whitefly adults were released to a single leaf encircled by a leaf cage (diameter, 45 mm; height, 30 mm). All the eggs on the *Arabidopsis* leaves were counted with a microscope after 10 d, and the number of eggs deposited per female was determined. Eight biological replicates were conducted in this experiment.

For the whitefly development experiment, 16 female adults were inoculated to a single leaf encircled by a leaf cage. After 2 d of oviposition, all adults were removed, and the eggs were allowed to develop. All pupae on the *Arabidopsis* leaves were counted with a microscope after 22 d, and the number of pupae per female was determined. Eight biological replicates were conducted in this experiment.

### Yeast two-hybrid analysis

The *Arabidopsis* Mate and Plate Library were used (Clontech, 630487). Full-length protein for βC1 was cloned into the pGBT9 vector to generate BD-βC1 construct. This was then used to screen against the full yeast library via the yeast mating system following the manufacturer’s protocol (Matchmaker Gold Yeast Two-Hybrid System, Clontech). To further confirm the interaction between βC1 and AtPIFs, full-length of *Arabidopsis* PIFs was cloned into the pGAD424 vector through LR reaction to generate AD-AtPIFs. The yeast strain Y2HGold was co-transformed with BD-βC1 and AD-AtPIF1/PIF3/PIF4/PIF5 constructs and plated on SD-Leu-Trp selective dropout medium. Colonies were transferred onto SD-Leu-Trp-His plates to verify positive clones. The empty vectors pGBT9 and pGAD424 were used as negative controls.

### Bimolecular fluorescence complementation (BiFC)

Fluorescence was observed owing to complementation of the βC1 fused with the C-terminal part of EYFP with one of PIFs fused with the N-terminal part of EYFP. Unfused nEYFP was used as a negative control. Leaves of 3-week-old *N. benthamiana* plants were infiltrated with *Agrobacterial* cells containing the constructs designed for this experiment. Two days after infiltration, fluorescence and DAPI staining were observed by confocal microscopy. Three independent plants were tested in one experiment. The experiment was repeated twice with similar results.

### Co-immunoprecipitation (Co-IP) assay

*A. tumefaciens* strains containing expression vectors of *35S:YFP* and *35S:AtPIF3-HA*, *35S:YFP-βC1* and *35S:AtPIF3-HA*, or *35S:YFP-βC1* and *35S:AtMYC2-HA* were co-injected into 3-week-old *N. benthamiana* leaf cells. YFP was used as negative control, and AtMYC2-HA was used as positive control. After infiltration, plants were maintained in the dark (in order to stabilize PIFs) for 2 d before protein extraction [52]. Total proteins were extracted from infiltrated leaf patches in 1 ml lysis buffer [50 mM Tris-HCl pH7.4, 150 mM NaCl, 2 mM MgCl_2_, 10% glycerol, 0.5% NP-40, 1 mM DTT, protease inhibitor cocktail (Roche, 32147600)]. Fifty milligram protein extracts were taken as input, and then the rest extracts were incubated with the GFP-Trap beads (ChromoTek, gta-20) for 1.5 h at 4°C. Immunoblotting was performed with anti-HA and anti-GFP antibodies (TransGen Biotech, HT801-02).

### Pull-down protein competitive interaction assay

The GST- and MBP-fusion proteins were separately purified using Glutathione sepharose (GE Healthcare, 17-5132-01) and Amylose resin (New England Biolabs, E8021S) beads as according to the manufacturer’s instructions. His-βC1 fusion proteins were purified using Ni-nitrilotriacetate (Ni-NTA) agarose (Qiagen, 30210) according to the manufacturer’s instructions. Indicated amounts of GST or His-βC1 were mixed with 2 μg of MBP-fusion proteins and 50 μL of Amylose resin overnight. After two washes with binding buffer (50 mM Tris-HCl, pH 7.5, 100 mM NaCl, 35 mM β-mercaptoethanol and 0.25% Triton X-100), 2 μg of GST-fusion proteins were added and the mixture was incubated for 3 h at 4°C. Beads were washed 6 times with binding buffer. The associated proteins were separated on 8 % SDS-polyacrylamide gels and detected by immunoblots using anti-GST antibody (TransGen Biotech, HT601-02).

### Quantitative RT-PCR

Total RNA was isolated using the RNeasy Plant Mini Kit (Qiagen, 74904), and 2000 ng of total RNA for each sample was reverse transcribed using the TransScript One-Step gDNA Removal and cDNA Synthesis SuperMix (TRAN, AT311-03). Three independent biological samples, each from an independent plant, were collected and analyzed. RT-qPCR was performed on the CFX 96 system (Bio-Rad) using Thunderbird SYBR qPCR mix (TOYOBO, QPS-201). The primers used for mRNA detection of target genes by real-time PCR are listed in Table S1. The *Arabidopsis Actin2* (At3g18780) mRNA was used as internal control.

### ChIP assay

Transgenic *Arabidopsis* plants expressing *35S:YFP-AtPIF3* and wild-type control Col-0 were used for ChIP assays. *Arabidopsis* seedlings were grown on MS medium for 12 days. 2.5 g of seedlings were harvested and fixed in 37 ml 1% formaldehyde solution under a vacuum for 10 min. Glycine was added to a final concentration of 0.125 M, and the sample was vacuum treated for an additional 5 min. After three washes with distilled water, samples were frozen in liquid nitrogen. ChIP experiments were performed as described using anti-GFP agarose beads (GFP track, gta-20) for immunoprecipitation [53]. The resulting DNA samples were purified with the QIA quick PCR purification kit (Qiagen, 28106). DNA fragments were analyzed by quantitative PCR, with the *Arabidopsis ACTIN2* (At3g18780) promoter as a reference. Enrichments were referred to the *35S:YFP-AtPIF3* against wild-type Col-0 seedlings. Primers of ChIP assays are listed in Table S1. The experiments were repeated with four independent biological samples, each from independent plants.

### Luciferase activity assay

*AtTPS10* Promoter: *luciferase* was used as a reporter construct. *35S:YFP*, *35S:AtPIF1*, *35S:AtPIF3*, *35S:AtPIF4*, *35S:AtPIF5*, *35S:AtMYC2* and *35S:βC1* were used as effector constructs. Nb leaves were agro-infiltrated with the constructs indicated in each figures. Two days after infiltration, leaves were harvested and the luciferase (LUC) activity of infiltrated leaf cells was quantified by microplate reader as described [22]. Each treatment was repeated eight times in one experiment.

### Protein extraction and western blot

For βC1 stability assays, construct containing *35S:myc-βC1* was infiltrated with *A. tumefaciens* strains (EHA105) and transiently expressed in leaves of four-week-old *N. benthamiana*. Plant samples were placed under different light conditions as indicated as in Figure 2. Total proteins were extracted from infiltrated leaf patches in 1 ml 2×NuPAGE LDS sample buffer (Invitrogen, NP0008) containing 0.05mL/mL β-mercaptoethanol, and protease inhibitor cocktail. Ten milligram protein extracts were taken for immunoblotting with anti-myc antibody (TransGen Biotech, HT101-01).

### Data analysis

Differences in whitefly performance, gene expression levels and average numbers of EYFP fluorescence were determined using Student’s *t*-tests for comparing two treatments or two lines. Differences in relative enrichment fold of DNA fragments in the promoter and relative LUC activity were determined using One-way ANOVA, followed by Duncan’s multiple range test for significant differences among different lines or different treatments. Differences in whitefly two-choice between different lines were analyzed by Wilcoxon matched pairs tests (with two dependent samples). All tests were carried out with GraphPad Prism.

### Accession numbers

Sequence data from this work can be found in Genebank/EMBL or The *Arabidopsis* Information Resource (www.Arabidopsis.org) under the following accession numbers: AtPIF1 (AT2G20180), AtPIF3 (AT1G09530), AtPIF4 (AT2G43010), AtPIF5 (AT3G59060), AtMYC2 (At1G32640), AtTPS10 (At2G24210), AtTPS14 (AT1G61680), AtTPS21 (AT5G23960), TYLCCNV βC1 (AJ421621).

## Acknowledgments

We thank Prof. Peter Quail for providing *pifq* mutant (University of California, Berkeley) and Dr. Jang In-Cheol (Temasek Life Sciences Laboratory, Singapore) for useful discussion.

## Author contributions

**Conceptualization**: Rongxiang Fang, Jian Ye.

**Data curation:** Pingzhi Zhao, Xuan Zhang, Yuqing Gong.

**Funding acquisition:** Jian Ye.

**Investigation:** Pingzhi Zhao, Xuan Zhang, Yuqing Gong.

**Methodology:** Pingzhi Zhao, Xuan Zhang, Yuqing Gong, Ning Wang.

**Project administration:** Jian Ye.

**Resources:** Dongqing Xu, Yanwei Sun.

**Supervision:** Jian Ye.

**Writing-original draft:** Pingzhi Zhao, Xuan Zhang, Yuqing Gong, Jian Ye.

**Writing-contributions:** Shu-Sheng Liu, Xing-Wang Deng, Daniel J. Kliebenstein, XuePing Zhou.

**Writing-review & editing:** Jian Ye.

## Supporting information

**S1 Fig.**
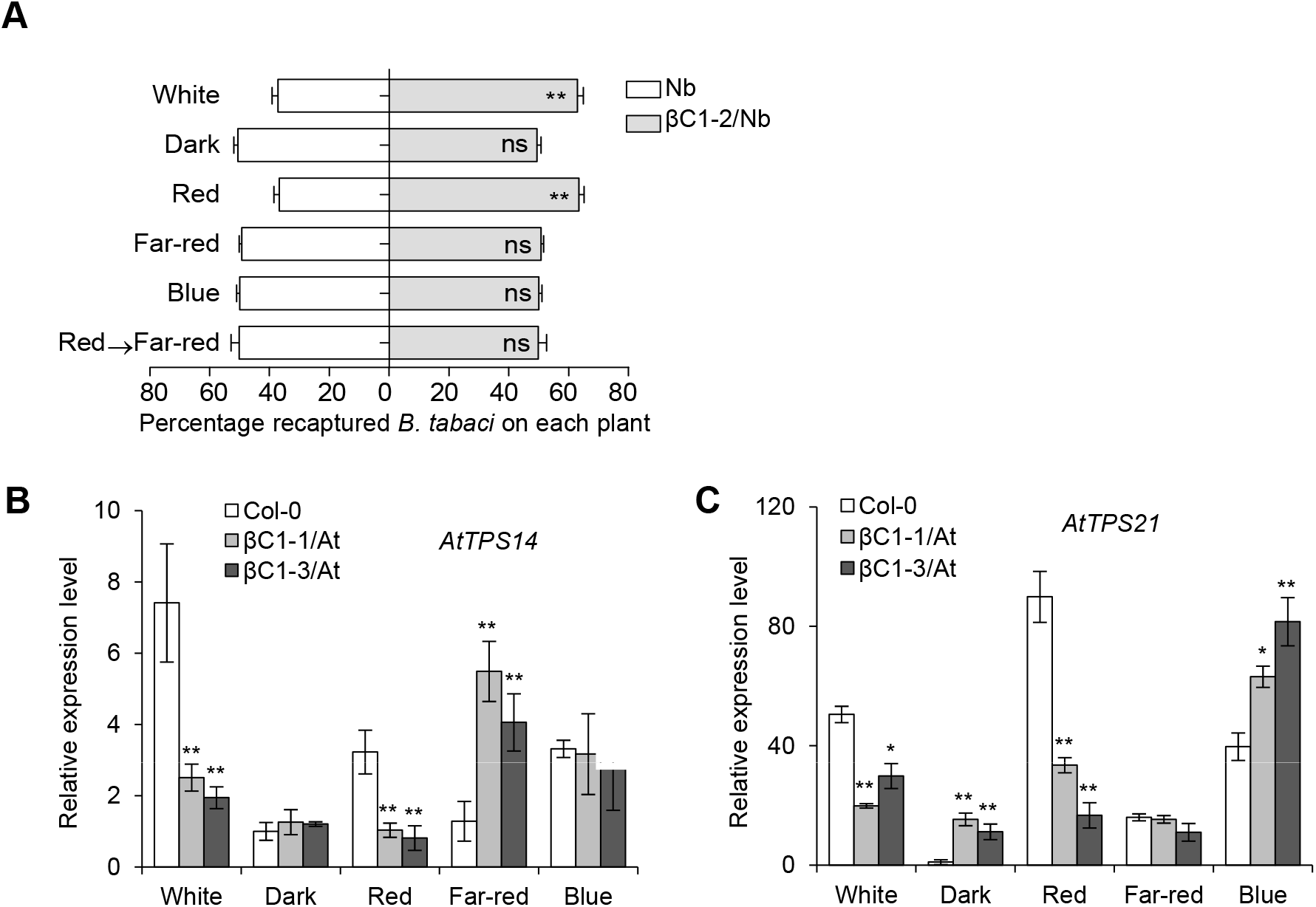
Begomovirus encodes βC1 to modulate light-regulated plant defense. **(A)** Whitefly preference on wild-type Nb and βC1 transgenic Nb plants (βC1-2/Nb) in response to white, dark, red, far-red, and blue light. Plants were placed under darkness for 24 h, followed by a 2 h light exposure and then performed whitefly choice experiments. Red→Far-red indicates that plants were firstly kept in darkness for 24 h, followed by a 2 h red light exposure, and then transferred to far-red light for 2 h. Values are mean + SD (n=6). Asterisks indicate significant differences between different treatments or lines (**, P< 0.01; ns, no significant differences; the Wilcoxon matched pairs test). **(B-C)** Relative expression levels of *AtTPS14* (B), and *AtTPS21* (C) in Col-0 and two βC1/At plants (βC1-1/At and βC1-3/At) under different light conditions. Values are mean ± SD (n=3) (*, P< 0.05; **, P< 0.01; Student’s *t*-test). The light was supplied by LED light sources, with irradiance fluency rates of: white (80 μmol m^−2^ sec^−1^), blue (15 μmol m^−2^ sec^−1^), red (20 μmol m^−2^ sec^−1^), and far-red (2 μmol m^−2^ sec^−1^).

**S2 Fig.**
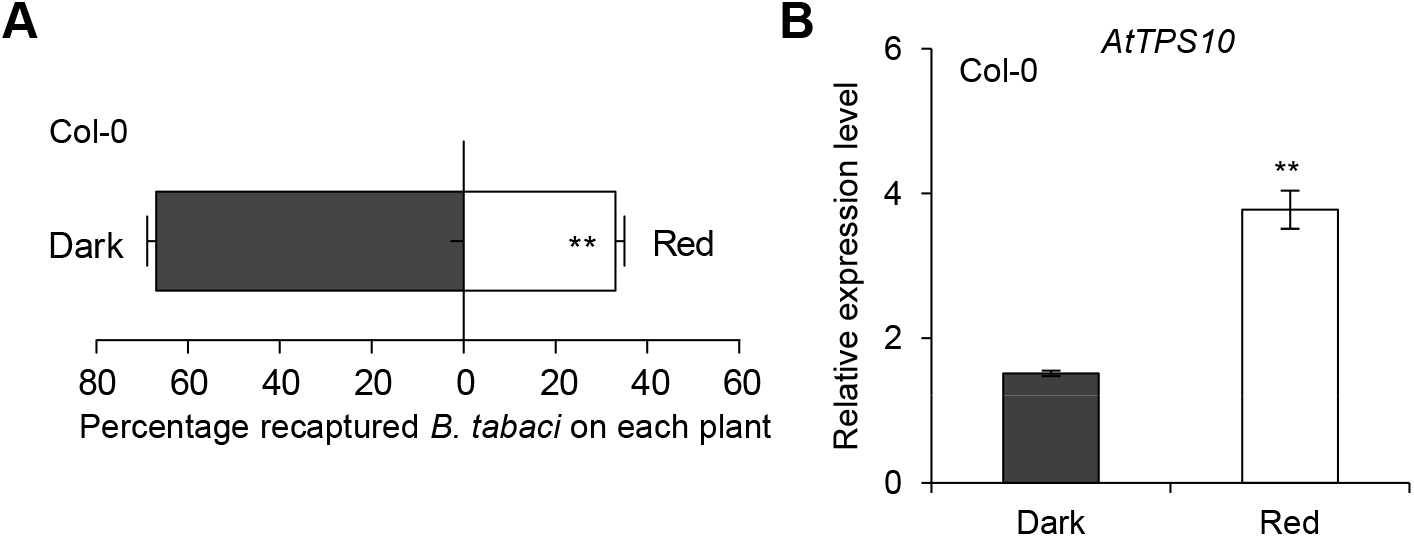
Red light plays a crucial role for plant defense against whitefly. **(A)** Whitefly preference (as percentage recaptured whiteflies out of 200 released) on wild-type Col-0 in response to darkness or red light. The plants were placed in darkness for 24 h prior to the 2 h dark or 2 h red light (20 μmol m^−2^ sec^−1^) treatments. Values are mean + SD (n=6). Asterisks indicate significant differences of whitefly preference between treatments (**, P< 0.01; the Wilcoxon matched pairs test). **(B)** Relative expression levels of *AtTPS10* in Col-0 plants exposed to darkness or red light. Values are mean ± SD (n=3). Asterisks indicate significant differences of *AtTPS10* expression in Col-0 plants between under darkness and red light (**, P< 0.01; Student’s *t*-test).

**S3 Fig.**
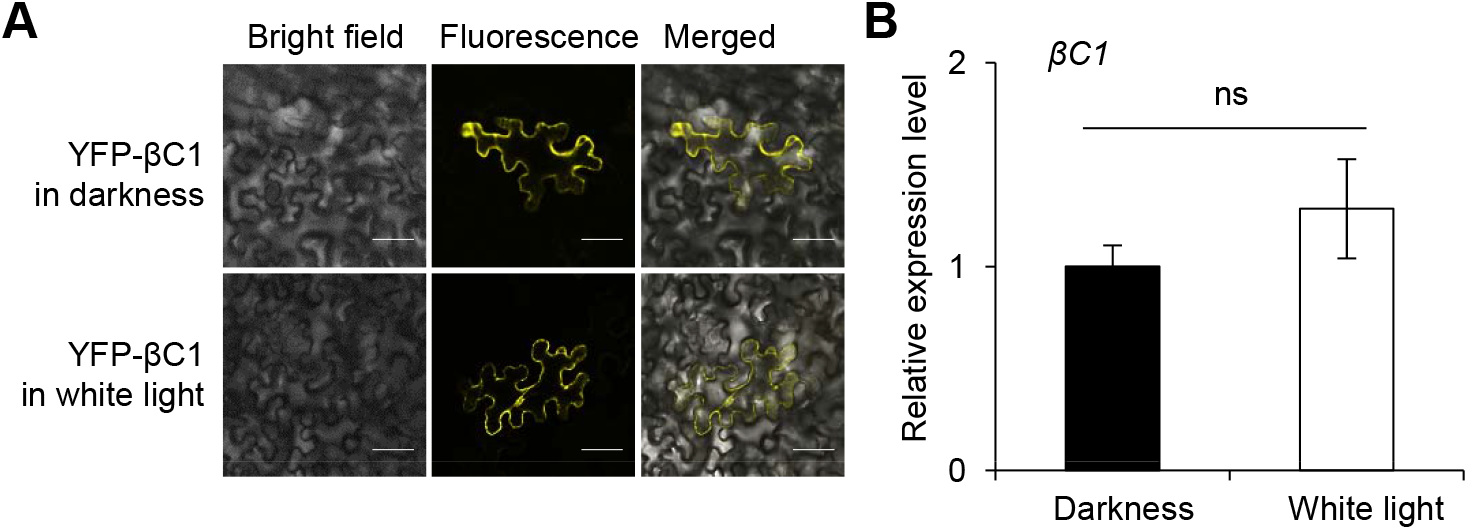
Light has no visible effect on the subcellular localization of βC1 protein or its transcript levels. **(A)** Subcellular localization of YFP-βC1 in *N. benthamiana* under darkness or white light condition. After transient inoculation of *35S:YFP-βC1*, plants were placed in the dark or in the white light for 48 h prior to the observation. Scale bars = 50 μm. **(B)** Relative expression levels of *βC1* in Col-0 plants in response to dark or white light. Values are means ± SD (n=3). ‘ns’ indicates no significant differences.

**S4 Fig.**
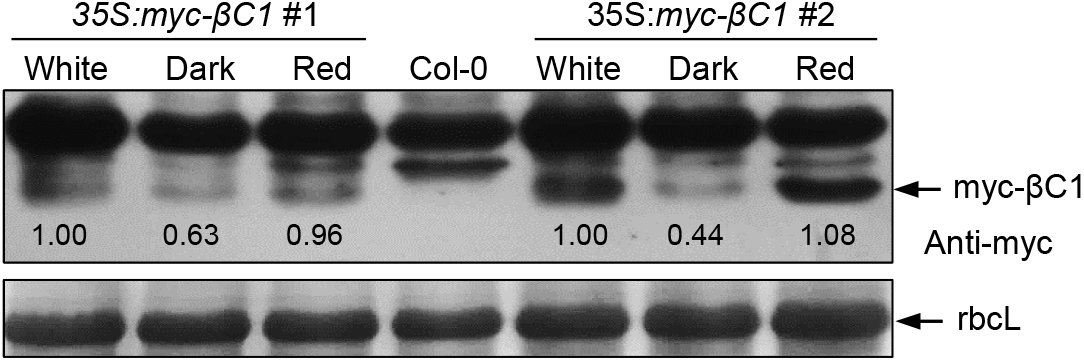
Red light promotes the stability of βC1 protein. Accumulation of βC1 proteins in two stable transgenic lines (*35S:myc-βC1* #1 and #2). Plants were placed under darkness for 24 h, followed by a 2 h light exposure. Samples were detected by immunoblot analysis using anti-myc antibody. Stained membrane bands of the large subunit of Rubisco (rbcL) were used as a loading control.

**S5 Fig.**
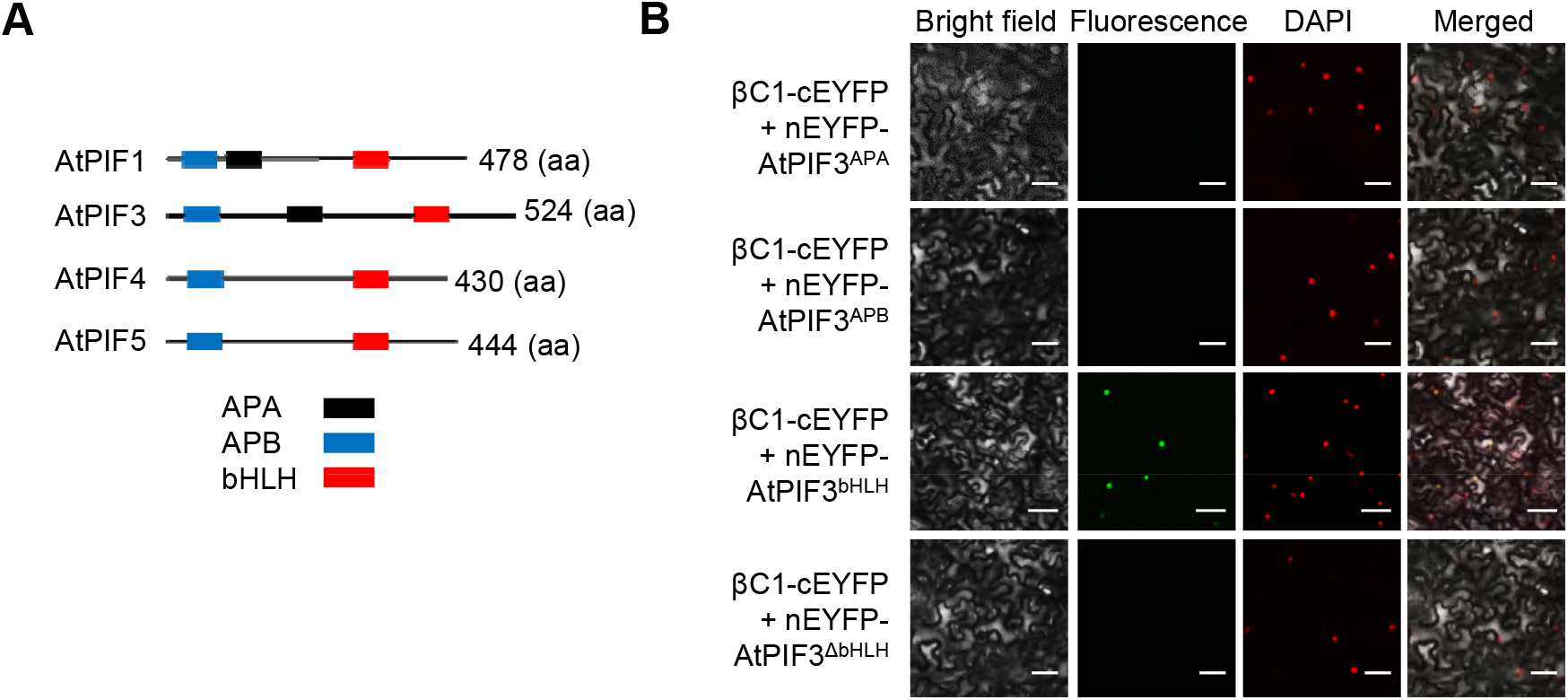
βC1 interacts with bHLH domain of AtPIFs protein. **(A)** Domain structure of AtPIFs proteins. Schematic diagrams of the AtPIFs polypeptide show the location of the consensus basic helix-loop-helix (bHLH) domain, which defines this transcription factor family, as well as the Active Phytochrome A-binding (APA) region and the Active Phytochrome B-binding (APB) region. **(B)** BiFC analysis of AtPIF3 derivative interaction with βC1 protein. The EYFP fluorescences were only observed owing to complementation of βC1-cEYFP with nEYFP-AtPIF3^bHLH^ in normal light. ∆bHLH indicates deletion of bHLH domain in AtPIF3. Scale bars = 50 μm.

**S6 Fig.**
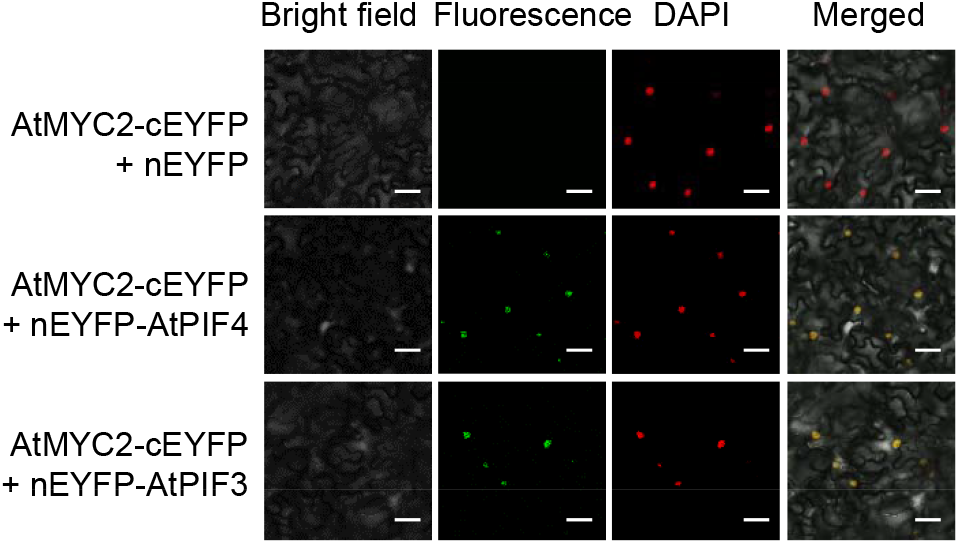
AtPIF proteins interact with MYC2. *In vivo* BiFC analysis of AtMYC2 interaction with AtPIFs (AtPIF3 or AtPIF4) in normal light. Scale bars = 50 μm.

**S7 Fig.**
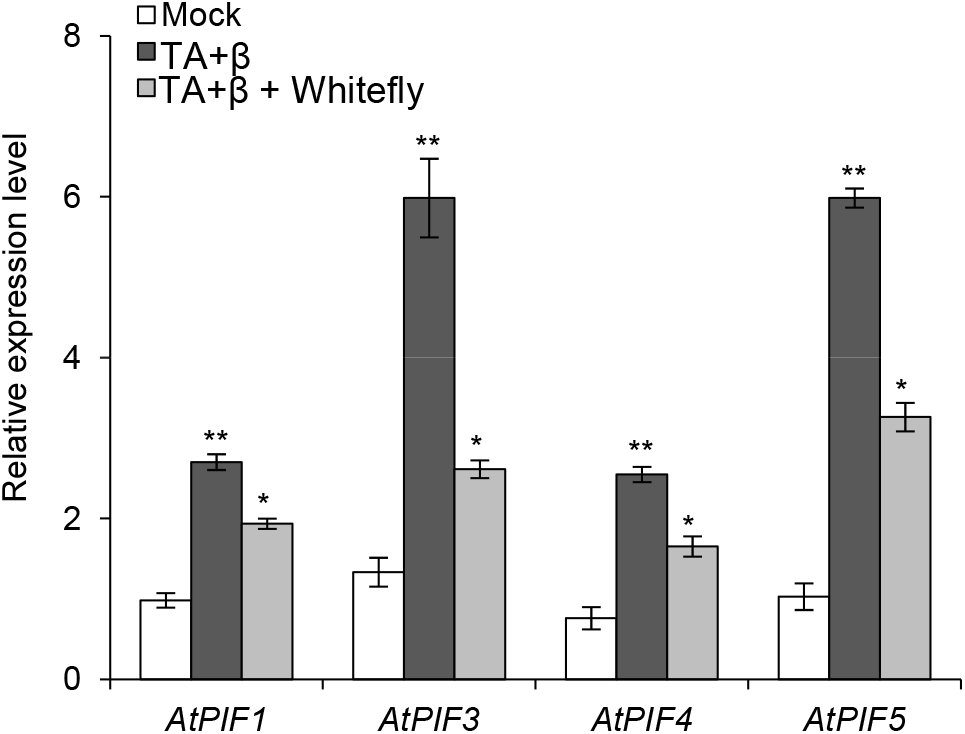
Begomovirus infection triggers PIFs transcription in *Arabidopsis*. Relative expression levels of *AtPIFs* in *Arabidopsis* plants. *Arabidopsis* Col-0 plants agroinfiltrated with the infectious clones of TA+β complex at 14 dpi, followed by infestation by whiteflies for 6 h. Total plant RNAs were extracted for qRT-PCR analysis. Uninfected Col-0 plants were used as mock. Values are means ± SD (n=3). Asterisks indicate significant differences of *AtPIF* genes expression between mock and infected-Col-0 plants (*, P< 0.05; **, P< 0.01; Student’s *t*-test).

**S8 Fig.**
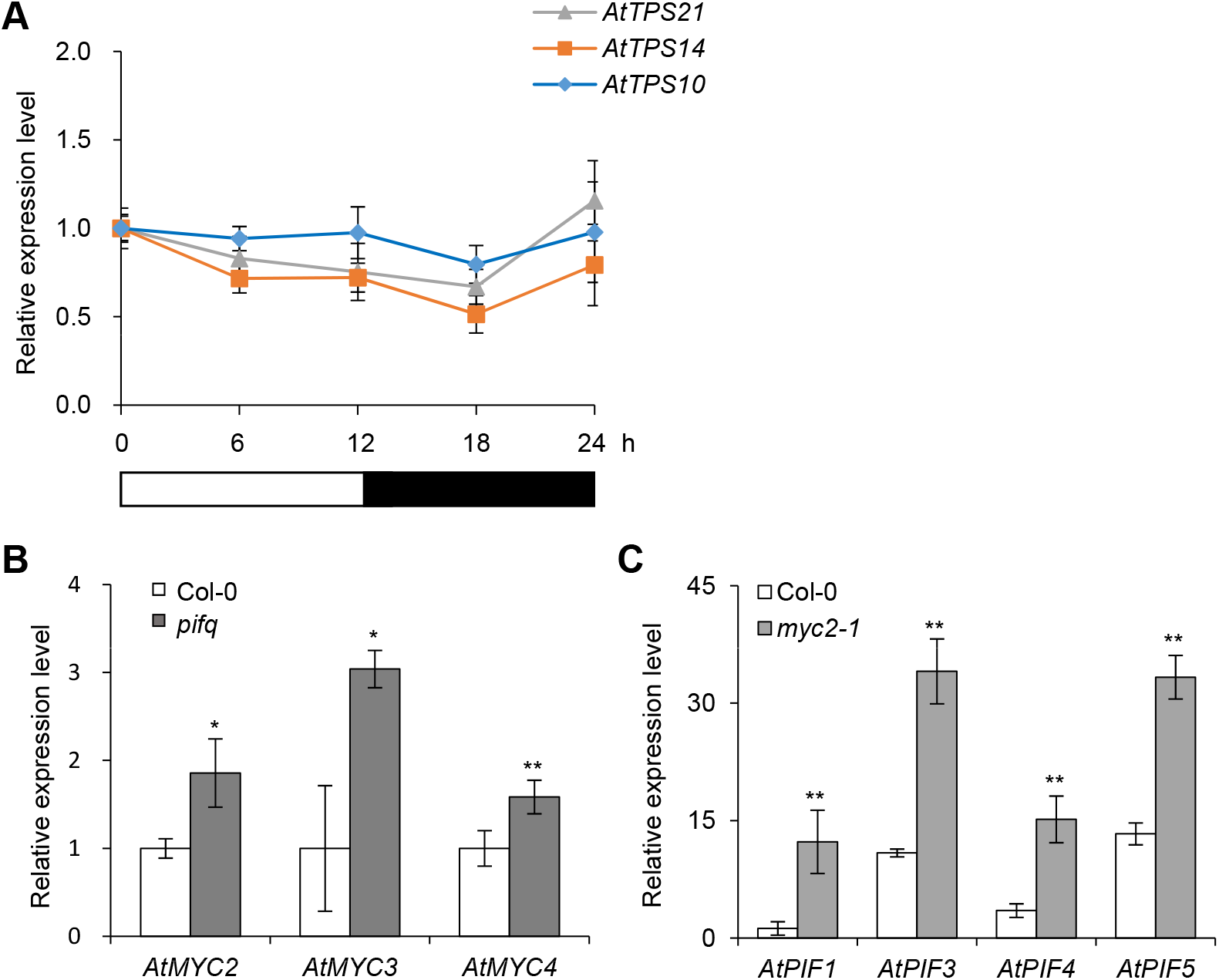
The expression of *PIFs* and *MYCs* is complemented and balanced regulation. **(A)** The expression pattern of *Arabidopsis TPS10/TPS14/TPS21* is constant during night and day time. Relative expression levels of *AtTPS* genes in Col-0 under 12 h light/12 h darkness. Values are mean ± SD (n=3). **(B)** Relative expression levels of *AtMYC* genes in Col-0 and *pifq* mutant plants under light. **(C)** Relative expression levels of *AtPIF* genes in Col-0 and *myc2-1* mutant plants under light. Values are mean ± SD (n=3). In figure **B**-**C**, asterisks indicate significant differences of genes expression between Col-0 and mutant plants (*, P< 0.05; **, P< 0.01; Student’s *t*-test).

**S9 Fig.**
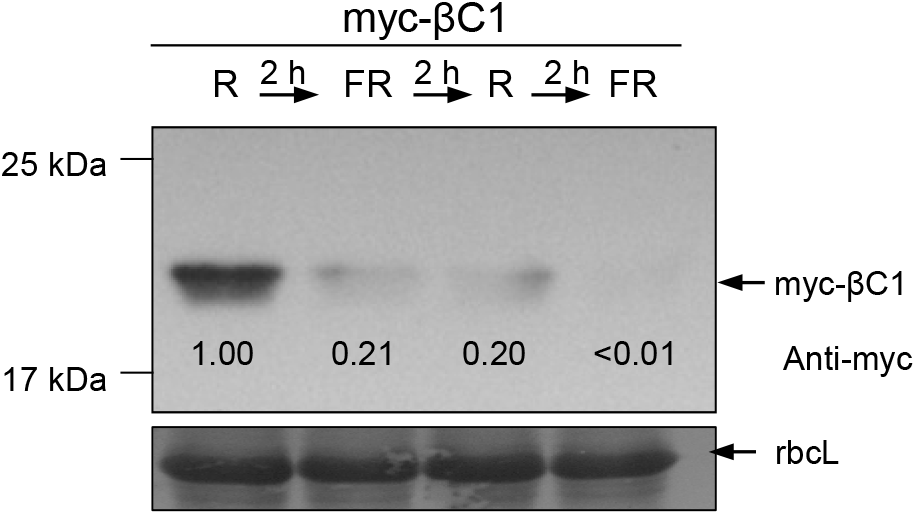
βC1 protein accumulation in continuous red light and far-red light. Accumulation of βC1 proteins in Nb plants after treated with continuous red light and far-red light. Plants were placed under darkness for 60 h, then transferred to continuous red light and far-red light for 2 h respectively. Samples were detected by immunoblot analysis using anti-myc antibody. Stained membrane bands of the large subunit of Rubisco (rbcL) were used as a loading control.

**S10 Fig.**
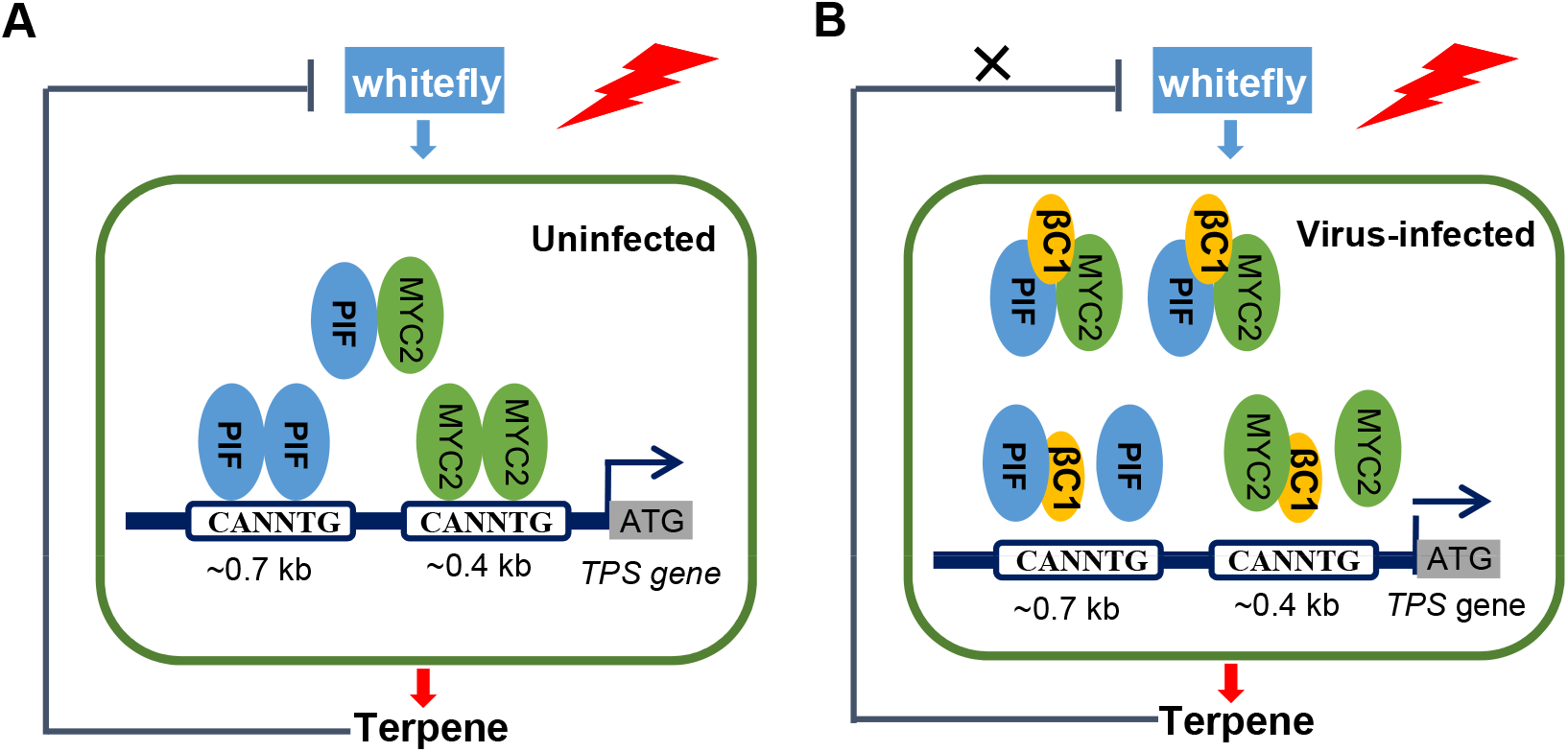
A working model of red-light regulated begomovirus-whitefly mutualism. **(A)** In uninfected plant, both plant PIFs and MYC2 mediate the transcription of *TPS* genes by respectively binding to different G-box-like elements of the promoter region, and activate *TPSs* transcription. Thus, red-light signal and JA signal fine-tune transcription of *TPS* genes in plants to defend against whitefly. **(B)** In begomovirus-infected plants, βC1 interacts with PIFs and MYC2, and inhibits their transcriptional activity by interfering with their homodimerization and promoting AtPIFs-AtMYC2 heterodimerization. Finally, the decreased terpene synthesis and in turn enhanced whitefly performance increase the probability of pathogen transmission.

**S1 Table.**
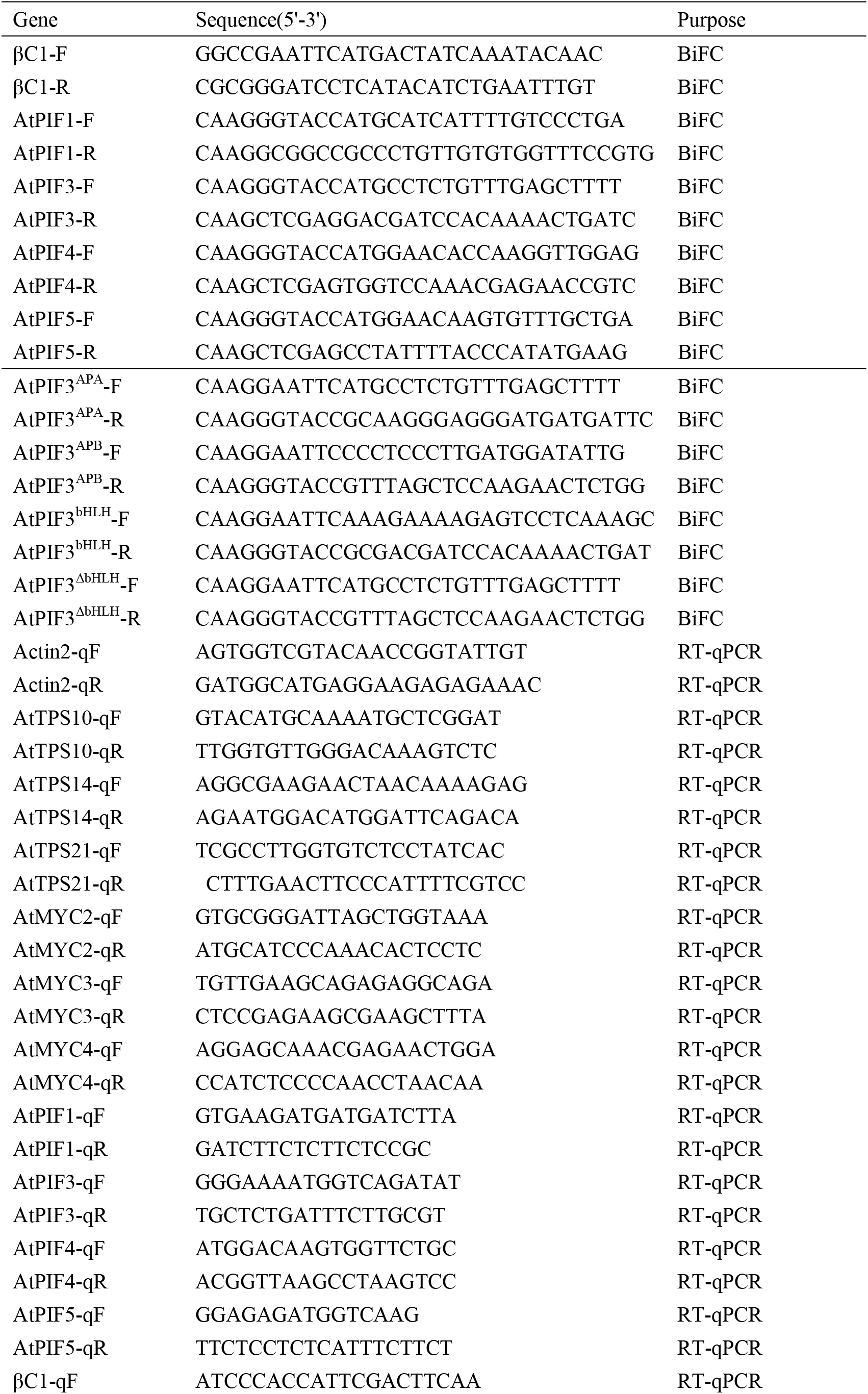
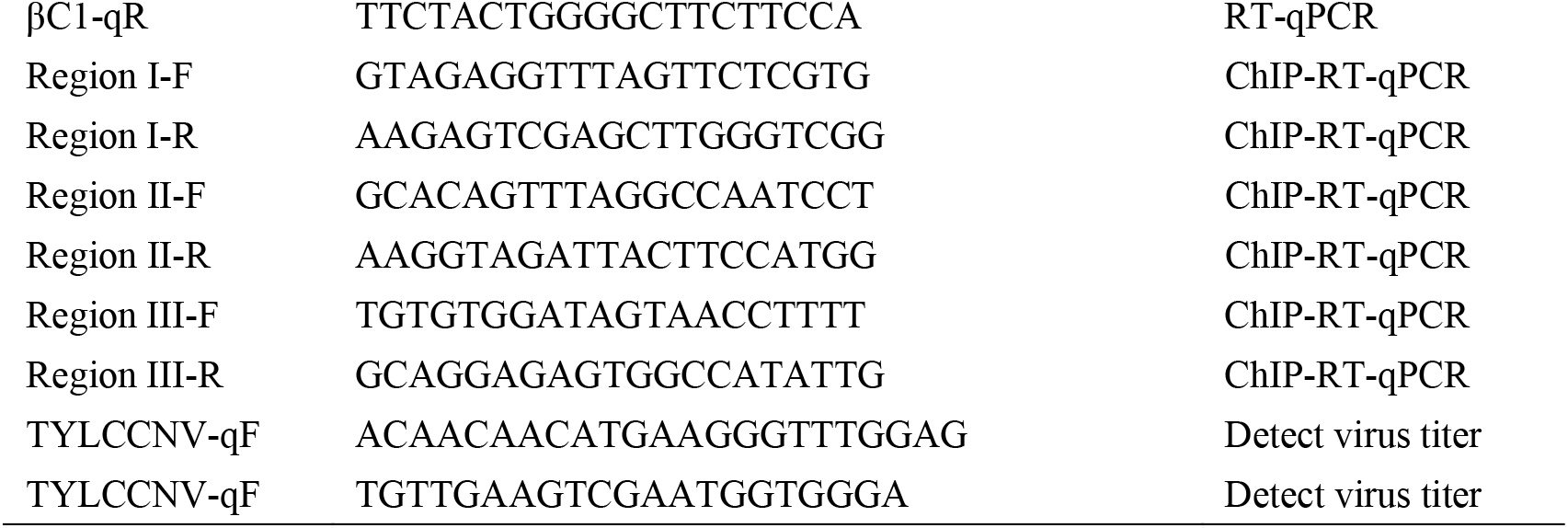
DNA primers used in this study.

## References

1. Deutsch CA, Tewksbury JJ, Tigchelaar M, Battisti DS, Merrill SC, Huey RB, and Naylor RL. Increase in crop losses to insect pests in a warming climate. Science 2018; 361(6405): 916–919. https://doi.org/10.1126/science.aat3466. PMID: 30166490.

2. Watts N, Amann M, Arnell N, Ayeb-Karlsson S, Belesova K, Boykoff M, Byass P, Cai W, Campbell-Lendrum, D, Capstick, S., et al. The 2019 report of The Lancet Countdown on health and climate change: ensuring that the health of a child born today is not defined by a changing climate. Lancet. 2019; 394(10211):1836–1878. https://doi.org/10.1016/S0140-6736(19)32596-6. PMID: 31733927.

3. Shah AA, Dillon ME, Hotaling S, and Woods HA. High elevation insect communities face shifting ecological and evolutionary landscapes. Curr Opin Insect Sci. 2020; 41:1–6. https://doi.org/10.1016/j.cois.2020.04.002. PMID: 32553896.

4. Navas-Castillo J, Fiallo-Olivé E, and Sánchez-Campos S. Emerging virus diseases transmitted by whiteflies. Annu Rev Phytopathol. 2011; 49: 219–248. https://doi.org/10.1146/annurev-phyto-072910-095235. PMID: 21568700.

5. Ronald R, and Beard CB. Vector-borne infections. Emerg Infect Dis. 2011; 17(5):769–770. https://doi.org/10.3201/eid1705.110310. PMID: 21529382.

6. Schuman MC, and Baldwin IT. Field studies reveal functions of chemical mediators in plant interactions. Chem Soc Rev. 2018; 47(14):5338–5353. https://doi.org/10.1039/c7cs00749c. PMID: 29770376.

7. Turlings TCJ, and Erb M. Tritrophic Interactions mediated by herbivore-induced plant volatiles: mechanisms, ecological relevance, and application potential. Annu Rev Entomol 2018; 63:433–452. https://doi.org/10.1146/annurev-ento-020117-043507. PMID: 29324043.

8. Ghanim M. (2014). A review of the mechanisms and components that determine the transmission efficiency of Tomato yellow leaf curl virus (Geminiviridae; Begomovirus) by its whitefly vector. Virus Res. 2014; 186:47–54. https://doi.org/10.1016/j.virusres. PMID: 24508344.

9. Eigenbrode SD, Bosque-Pérez NA, and Davis TS. Insect-borne plant pathogens and their vectors: ecology, evolution, and complex interactions. Annu Rev Entomol. 2018; 63:169–191. https://doi.org/10.1146/annurev-ento-020117-043119. PMID: 28968147.

10. Zhao P, Yao X, Cai C, Li R, Du J, Sun Y, Wang M, Zou Z, Wang Q, Kliebenstein, D.J., et al. Viruses mobilize plant immunity to deter nonvector insect herbivores. Sci Adv. 2019; 5(8):eaav9801. https://doi.org/10.1126/sciadv.aav9801. PMID: 31457079.

11. Scheres, B., and van der Putten, W.H. The plant perceptron connects environment to development. Nature, 2017;543(7645), 337–345. https://doi.org/10.1038/nature22010. PMID: 28300110.

12. Stam, J.M., Kroes, A., Li, Y., Gols, R., van Loon, J.J.A., Poelman, E.H., and Dicke, M. Plant interactions with multiple insect herbivores: from community to genes. Annu Rev Plant Biol. 2014; 65, 689–713. https://doi.org/10.1146/annurev-arplant-050213-035937. PMID: 24313843

13. Schuman, M.C., and Baldwin, I.T. The layers of plant responses to insect herbivores. Annu Rev Entomol. 2016; 61, 373–394. https://doi.org/10.1146/annurev-ento-010715-023851. PMID: 26651543

14. Ballaré, C.L. Light regulation of plant defense. Annu Rev Plant Biol. 2014; 65, 335–363. https://doi.org/10.1146/annurev-arplant-050213-040145. PMID: 24471835

15. Douma, J.C., de Vries, J., Poelman, E.H., Dicke, M., Anten, N.P.R., and Evers, J.B. Ecological significance of light quality in optimizing plant defence. Plant Cell Environ. 2019;42(3), 1065–1077. https://doi.org/10.1111/pce.13524. PMID: 30702750

16. Pham, V.N., Kathare, P.K., and Huq, E. Phytochromes and phytochrome interacting factors. Plant Physiol. 2018; 176 (2): 1025. https://doi.org/doi.org/10.1104/pp.17.01384. PMID: 29138351

17. Ni, M., Tepperman, J.M., and Quail, P.H. PIF3, a phytochrome-interacting factor necessary for normal photoinduced signal transduction, is a novel basic helix-loop-helix protein. Cell, 1998 2998; 95(5): 657–667. https://doi.org/10.1016/s0092-8674(00)81636-0. PMID: 9845368

18. Toledo-Ortiz, G., Huq, E., and Quail, P.H. The Arabidopsis basic/helix-loop-helix transcription factor family. The Plant Cell, 2003; 15(8): 1749–1770. https://doi.org/10.1105/tpc.013839. PMID: 12897250

19. Leivar, P., and Quail, P.H. PIFs: pivotal components in a cellular signaling hub. Trends Plant Sci. 2011; 16(1):19–28. https://doi.org/10.1016/j.tplants.2010.08.003. PMID: 20833098

20. Dicke, M., and Baldwin, I.T. The evolutionary context for herbivore-induced plant volatiles: beyond the ‘cry for help’. Trends Plant Sci. 2010; 15(3):167–175. https://doi.org/10.1016/j.tplants.2009.12.002. PMID: 20047849

21. Hong, G.J., Xue, X.Y., Mao, Y.B., Wang, L.J., and Chen, X.Y. Arabidopsis MYC2 interacts with DELLA proteins in regulating sesquiterpene synthase gene expression. The Plant Cell, 2012; 24(6): 2635–2648. https://doi.org/10.1105/tpc.112.098749. PMID: 22669881

22. Li, R., Weldegergis, B.T., Li, J., Jung, C., Qu, J., Sun, Y., Qian, H., Tee, C., van Loon, J.J., and Dicke, M. Virulence factors of geminivirus interact with MYC2 to subvert plant resistance and promote vector performance. The Plant Cell, 2014; 26(12): 4991–5008. https://doi.org/10.1105/tpc.114.133181. PMID: 25490915

23. Wu, X., Xu, S., Zhao, P., Zhang, X., Yao, X., Sun, Y., Fang, R., and Ye, J. The Orthotospovirus nonstructural protein NSs suppresses plant MYC-regulated jasmonate signaling leading to enhanced vector attraction and performance. PLoS Pathog. 2019; 15(6): e1007897. https://doi.org/10.1371/journal.ppat.1007897. PMID: 31206553

24. Wu, X., and Ye, J. Manipulation of jasmonate signaling by plant viruses and their insect vectors. Viruses, 2020; 12(2):148. https://doi.org/10.3390/v12020148. PMID: 32012772

25. Kegge, W., Weldegergis, B.T., Soler, R., Eijk, M.V.-V., Dicke, M., Voesenek, L.A.C.J., and Pierik, R. (2013). Canopy light cues affect emission of constitutive and methyl jasmonate-induced volatile organic compounds in Arabidopsis thaliana. New Phytol. 2013; 200(3):861–874. https://doi.org/10.1111/nph.12407. PMID: 23845065

26. Luan, J.B., Yao, D.M., Zhang, T., Walling, L.L., Yang, M., Wang, Y.J., and Liu, S.S. Suppression of terpenoid synthesis in plants by a virus promotes its mutualism with vectors. Ecol Lett. 2013; 16(3):390–398. https://doi.org/10.1111/ele.12055. PMID: 23279824

27. Zhou, X. Advances in understanding begomovirus satellites. Annu Rev Phytopathol. 2013; 51: 357–381. https://doi.org/10.1146/annurev-phyto-082712-102234. PMID: 23915133

28. Li, F., Yang, X., Bisaro, D.M., and Zhou, X. The βC1 Protein of geminivirus–betasatellite complexes: a target and repressor of host defenses. Mol Plant, 2018;11(12):1424–1426. https://doi.org/10.1016/j.molp.2018.10.007, PMID: 30404041

29. Yang, X., Guo, W., Li, F., Sunter, G., and Zhou, X. Geminivirus-associated betasatellites: exploiting chinks in the antiviral arsenal of plants. Trends Plant Sci. 2019;24(6): 519–529. https://doi.org/10.1016/j.tplants.2019.03.010, PMID: 31003895

30. Bleeker, P.M., Diergaarde, P.J., Ament, K., Guerra, J., Weidner, M., Schutz, S., de Both, M.T., Haring, M.A., and Schuurink, R.C. The role of specific tomato volatiles in tomato-whitefly interaction. Plant Physiol. 2009; 151(2): 925–935. https://doi.org/10.1104/pp.109.142661, PMID: 19692533

31. Shibuya, T., Komuro, J., Hirai, N., Sakamoto, Y., Endo, R., and Kitaya, Y. (2010). Preference of sweetpotato whitefly adults to cucumber seedlings grown under two different light sources. Horttechnol. 2010; 20(5):873–876. https://doi.org/10.21273/HORTTECH.20.5.873.

32. Briscoe, A.D., and Chittka, L. The evolution of color vision in insects. Annu Rev Entomol. 2001; 46: 471. https://doi.org/10.1146/annurev.ento.46.1.471. PMID: 11112177

33. Liu, S.S., De Barro, P.J., Xu, J., Luan, J.B., Zang, L.S., Ruan, Y.M., and Wan, F.H. Asymmetric mating interactions drive widespread invasion and displacement in a whitefly. Science, 2007; 318 (5857):1769–1772. https://doi.org/10.1126/science.1149887. PMID: 17991828

34. Paik, I., Kathare, P.K., Kim, J.I., and Huq, E. Expanding roles of PIFs in signal integration from multiple processes. Molecular Plant 2017;10(8): 1035–1046. https://doi.org/10.1016/j.molp.2017.07.002. PMID: 28711729

35. Martínez-García, J.F., Huq, E., and Quail, P.H. Direct targeting of light signals to a promoter element-bound transcription factor. Science, 2000; 288(5467):859–863. https://doi.org/10.1126/science.288.5467.859. PMID: 10797009

36. Zhang, Y., Mayba, O., Pfeiffer, A., Shi, H., Tepperman, J.M., Speed, T.P., and Quail, P.H. A quartet of PIF bHLH factors provides a transcriptionally centered signaling hub that regulates seedling morphogenesis through differential expression-patterning of shared target genes in Arabidopsis. PLoS Genet. 2013; 9(1), e1003244. https://doi.org/10.1371/journal.pgen.1003244. PMID: 23382695

37. Zhang, X., Ji, Y., Xue, C., Ma, H., Xi, Y., Huang, P., Wang, H., An, F., Li, B., Wang, Y., et al. Integrated regulation of apical hook development by transcriptional coupling of EIN3/EIL1 and PIFs in Arabidopsis. The Plant Cell, 2018;30(9):1971–1988. https://doi.org/10.1105/tpc.18.00018. PMID: 30104405

38. Velásquez, A.C., Castroverde, C.D.M., and He, S.Y. Plant–pathogen warfare under changing climate conditions. Cur Biol. 2018;28(10):R619–R634. https://doi.org/10.1016/j.cub.2018.03.054. PMID: 29787730

39. Carr, J.P., Murphy, A.M., Tungadi, T., and Yoon, J.-Y. Plant defense signals: Players and pawns in plant-virus-vector interactions. Plant Sci. 2019; 279:87–95. https://doi.org/10.1016/j.plantsci.2018.04.011. PMID: 30709497

40. Fereres, A., Peñaflor, M.F., Favaro, C.F., Azevedo, K.E., Landi, C.H., Maluta, N.K., Bento, J.M., and Lopes, J.R. Tomato infection by whitefly-transmitted circulative and non-circulative viruses induce contrasting changes in plant volatiles and vector behaviour. Viruses. 2016; 8(8):225. https://doi.org/10.3390/v8080225. PMID: 27529271

41. Mann, R.S., Ali, J.G., Hermann, S.L., Tiwari, S., Pelz-Stelinski, K.S., Alborn, H.T., and Stelinski, L.L. Induced release of a plant-defense volatile ‘Deceptively’ attracts insect vectors to plants infected with a bacterial pathogen. PLoS Pathog. 2012; 8(3): e1002610. https://doi.org/10.1371/journal.ppat.1002610. PMID: 22457628

42. Chico, J.M., Fernández-Barbero, G., Chini, A., Fernández-Calvo, P., Díez-Díaz, M., and Solano, R. Repression of jasmonate-dependent defenses by shade involves differential regulation of protein stability of MYC transcription factors and their JAZ repressors in Arabidopsis. The Plant Cell, 2014; 26(3):1967–1980. https://doi.org/10.1105/tpc.114.125047. PMID: 24824488

43. Castillon, A., Shen, H., and Huq, E. Phytochrome Interacting Factors: central players in phytochrome-mediated light signaling networks. Trends Plant Sci. 2007; 12(11):514–521. https://doi.org/10.1016/j.tplants.2007.10.001. PMID: 17933576

44. Lee, N., and Choi, G. Phytochrome-interacting factor from Arabidopsis to liverwort. Curr Opin Plant Biol. 2017; 35: 54–60. https://doi.org/10.1016/j.pbi.2016.11.004. PMID: 27875778

45. Franklin, K.A., and Quail, P.H. Phytochrome functions in Arabidopsis development. J Exp Bot. 2009; 61(1): 11–24. https://doi.org/10.1093/jxb/erp304. PMID: 19815685

46. Li, J., Li, G., Wang, H., and Wang Deng, X. Phytochrome signaling mechanisms. Arabidopsis Book. 2011(9): e0148. https://doi.org/10.1199/tab.0148. PMID: 22303272

47. Haxim, Y., Ismayil, A., Jia, Q., Wang, Y., Zheng, X., Chen, T., Qian, L., Liu, N., Wang, Y., Han, S., et al. Autophagy functions as an antiviral mechanism against geminiviruses in plants. eLife. 2017; 6: e23897. https://doi.org/10.7554/eLife.23897. PMID: 28244873

48. Yang, J.Y., Iwasaki, M., Machida, C., Machida, Y., Zhou, X., and Chua, N.H.βC1, the pathogenicity factor of TYLCCNV, interacts with AS1 to alter leaf development and suppress selective jasmonic acid responses. Genes Dev. 2008; 22(18): 2564–2577. https://doi.org/10.1101/gad.1682208. PMID: 18794352

49. Leivar, P., Tepperman, J.M., Monte, E., Calderon, R.H., Liu, T.L., and Quail, P.H. Definition of early transcriptional circuitry involved in light-induced reversal of PIF-imposed repression of photomorphogenesis in young Arabidopsis seedlings. The Plant Cell, 2009; 21(11):3535–3553. https://doi.org/10.1105/tpc.109.070672. PMID: 19920208

50. Lorenzo, O., Chico, J.M., Sánchez-Serrano, J.J., and Solano, R. JASMONATE-INSENSITIVE1 encodes a MYC transcription factor essential to discriminate between different jasmonate-regulated defense responses in Arabidopsis. The Plant Cell, 2004; 16(7):1938–1950. https://doi.org/10.1105/tpc.022319. PMID: 15208388

51. Cui, X., Tao, X., Xie, Y., Fauquet, C.M., and Zhou, X. A DNAbeta associated with Tomato yellow leaf curl China virus is required for symptom induction. J Virol. 2004; 78(24): 13966–13974. https://doi.org/10.1128/JVI.78.24.13966-13974.2004. PMID: 15564504

52. Quail, P.H. Phytochrome photosensory signalling networks. Nat Rev Mol Cell Biol. 2002; 3(2): 85–93. https://doi.org/10.1038/nrm728. PMID: 11836510

53. Ye, J., Yang, J., Sun, Y., Zhao, P., Gao, S., Jung, C., Qu, J., Fang, R., and Chua, N.H. Geminivirus activates ASYMMETRIC LEAVES 2 to accelerate cytoplasmic DCP2-mediated mRNA turnover and weakens RNA silencing in Arabidopsis. PLoS Pathog. 2015; 11(10):e1005196 https://doi.org/10.1371/journal.ppat.1005196. PMID: 26431425

